# Deep generative modeling of transcriptional dynamics for RNA velocity analysis in single cells

**DOI:** 10.1101/2022.08.12.503709

**Authors:** Adam Gayoso, Philipp Weiler, Mohammad Lotfollahi, Dominik Klein, Justin Hong, Aaron Streets, Fabian J. Theis, Nir Yosef

## Abstract

RNA velocity has been rapidly adopted to guide the interpretation of transcriptional dynamics in snapshot single-cell transcriptomics data. Current approaches for estimating and analyzing RNA velocity can empirically reveal complex dynamics but lack effective strategies for quantifying the uncertainty of the estimate and its overall applicability to the system of interest. Here, we present veloVI (velocity variational inference), a deep generative modeling framework for estimating RNA velocity. veloVI learns a gene-specific dynamical model of RNA metabolism and provides a transcriptome-wide quantification of velocity uncertainty. We show in a series of examples that veloVI compares favorably to previous approaches for inferring RNA velocity with improvements in fit to the data, consistency across transcriptionally similar cells, and stability across preprocessing pipelines for quantifying RNA abundance. Further, we demonstrate that properties unique to veloVI, such as posterior velocity uncertainty, can be used to assess the appropriateness of analysis with velocity to the data at hand. Finally, we highlight veloVI as a flexible framework for modeling transcriptional dynamics by adapting the underlying dynamical model to use time-dependent transcription rates.

## Introduction

Advances in single-cell RNA sequencing (scRNA-seq) technologies have facilitated the high-resolution dissection of the mechanisms underlying cellular differentiation and other temporal processes^1–3^. Although scRNA-seq is a destructive assay, a widely-used set of computational approaches leverage the asynchronous nature of dynamical biological processes to order cells along a so-called pseudotime in the task of trajectory inference^4–7^. Traditional methods for trajectory inference typically require the initial state of the underlying biological process to be known and use manifold learning to determine a metric space in which distances capture changes in differentiation state.

Recently, RNA velocity has emerged as a bottom-up, mechanistic approach for the trajectory inference task. RNA velocity, which describes the change of spliced messenger RNA (mRNA) over time, makes use of the observation that both unspliced and spliced RNA transcripts are simultaneously detected with standard scRNA-seq protocols^8^. Upon estimation, RNA velocity is typically incorporated into analyses in two ways: (1) to infer a cell-specific differentiation pseudotime, or (2) to construct a transition matrix inducing a Markov chain over the data that can be used to determine initial, transient, and terminal subpopulations of cells^9^.

There are currently two popular methods for estimating RNA velocity. The first, which we refer to as the *steady-state model*, assumes (1) constant rates of transcription and degradation of RNA, (2) a single, global splicing rate^8,10^, (3) that the cellular dynamics reached an equilibrium in the induction phase and do not include basal transcription, and (4) gene-wise independence. The second method, referred to as the *EM model*, we previously described and implemented in the scVelo package^11^. The *EM model* relaxes the assumption of the system having reached a steady-state, infers the full set of transcriptional parameters, and estimates a latent time per cell per gene by formulating the problem in an expectation-maximization (EM) framework.

While these approaches for estimating RNA velocity have been successfully used to interpret single-cell dynamics^12,13^, they also suffer from various limitations derived from their modeling assumptions as well as downstream usage^14–17^. For example, for both methods, there is no global notion of uncertainty. Thus, assessing the robustness of the RNA velocity estimate, or deciding to what extent velocity analysis is appropriate for a given dataset can be difficult. Although the *EM model* can be used to rank putative “driving” genes by their likelihood, there is no direct connection between gene likelihood, visualization, and correctness. For example, in the case of dentate gyrus neurogenesis, visualization of RNA velocity suggests that Granule mature cells develop into their immature counterparts even though a selection of genes with a high likelihood suggests the reverse (correct) dynamics^11^.

Estimation of RNA velocity with current approaches is also tightly coupled to the parameterization of the differential equations underlying transcription. Assumptions like constant transcription, splicing, and degradation rates may be too simple to explain dynamics that arise in multi-lineage^14^ or even single-lineage^18^ cell differentiation. The methods outlined to estimate RNA velocity lack extensibility and flexibility to adapt to these more complicated, real-world scenarios. Emerging technologies like VASA-seq^19^, which have greater sensitivity for unspliced RNA detection, may also provide sufficient signal to fit more complex models (e.g., going beyond two promoter states^20^); thus, a framework is necessary to rapidly test and update model assumptions.

To address these issues, we present veloVI (velocity variational inference), a deep generative model for estimation of RNA velocity. VeloVI reformulates the inference of RNA velocity via a model that shares information between all cells and genes while learning the same quantities, namely kinetic parameters and latent time, as in the *EM model*. This reformulation leverages advances in deep generative modeling^21^, which have become integral to many single-cell omics analytical tasks such as multi-modal data integration^22,23^, perturbation modeling^24,25^, and data correction^26^. As its output, veloVI returns an empirical posterior distribution of RNA velocity (matrix of cells by genes by posterior samples), which can be incorporated into the downstream analysis of the results. Here, we show that veloVI represents a significant improvement over the *EM model* in terms of fit to the data. Additionally, it provides a layer of interpretation and model criticism lacking from previous methods while also greatly improving flexibility for model extensions.

We use veloVI to enhance analyses of velocity at the level of cells, genes, and whole datasets. At the level of a cell, veloVI illuminates cell states that have directionality estimated with high uncertainty, which adds a notion of confidence to the velocity stream and highlights regions of the phenotypic manifold that warrant further investigation and more careful interpretation. We couple this analysis with a new metric called velocity coherence that explains the extent to which a gene agrees/disagrees with the inferred directionality. At the level of genes and datasets, we propose a novel permutation-based technique using veloVI that can identify partially-observed dynamics or systems in steady-states. This can be used to determine the extent to which RNA velocity analysis is suitable for a particular dataset.

Finally, veloVI is an extensible framework to fit more sophisticated transcriptional models. We highlight this flexibility by extending the current transcriptional model with a time-dependent transcription rate and show how this extension can improve the model fit.

## Results

### Variational inference for estimating RNA velocity

VeloVI posits that the unspliced and spliced abundances of RNA for each gene in a cell are generated as a function of kinetic parameters (transcription, splicing, degradation rates), a latent time, and a latent transcriptional state (i.e., induction state, repression state, and their respective steady states). Additionally, veloVI posits that the latent times for each gene in a cell are tied via a low-dimensional latent variable that we call the cell representation. These representations capture the notion that the observed state of a cell is a composition of multiple concomitant processes that together span the phenotypic manifold^1^. This modeling choice is justified by the observation that with the *EM model*, which is fit independently per gene, the inferred latent time matrix (of shape cells by genes) has a low-rank structure (but notably, not rank one; Supplementary Fig. 1).

The complete architecture of veloVI manifests as a variational autoencoder ^30^ with encoder and decoder components. The encoder neural networks take the unspliced and spliced abundances of a cell as input and output the posterior parameters for the cell representation and latent transcriptional state variables. The gene-wise, state-specific, latent time is parametrized by a neural network that takes a sample of the cell representation as input. The likelihood of cellular unspliced and spliced abundances is then a function of the latent time, the kinetic rate parameters, and the state assignment probabilities (Figure 1a; Methods). The model’s parameters are optimized simultaneously using standard gradient-based procedures. After optimization, the cell-gene-specific velocity is computed as a function of the degradation rate, the splicing rate, and the fitted unspliced and spliced abundances, which directly incorporate the posterior distributions over time and transcriptional state.

**Fig. 1.**
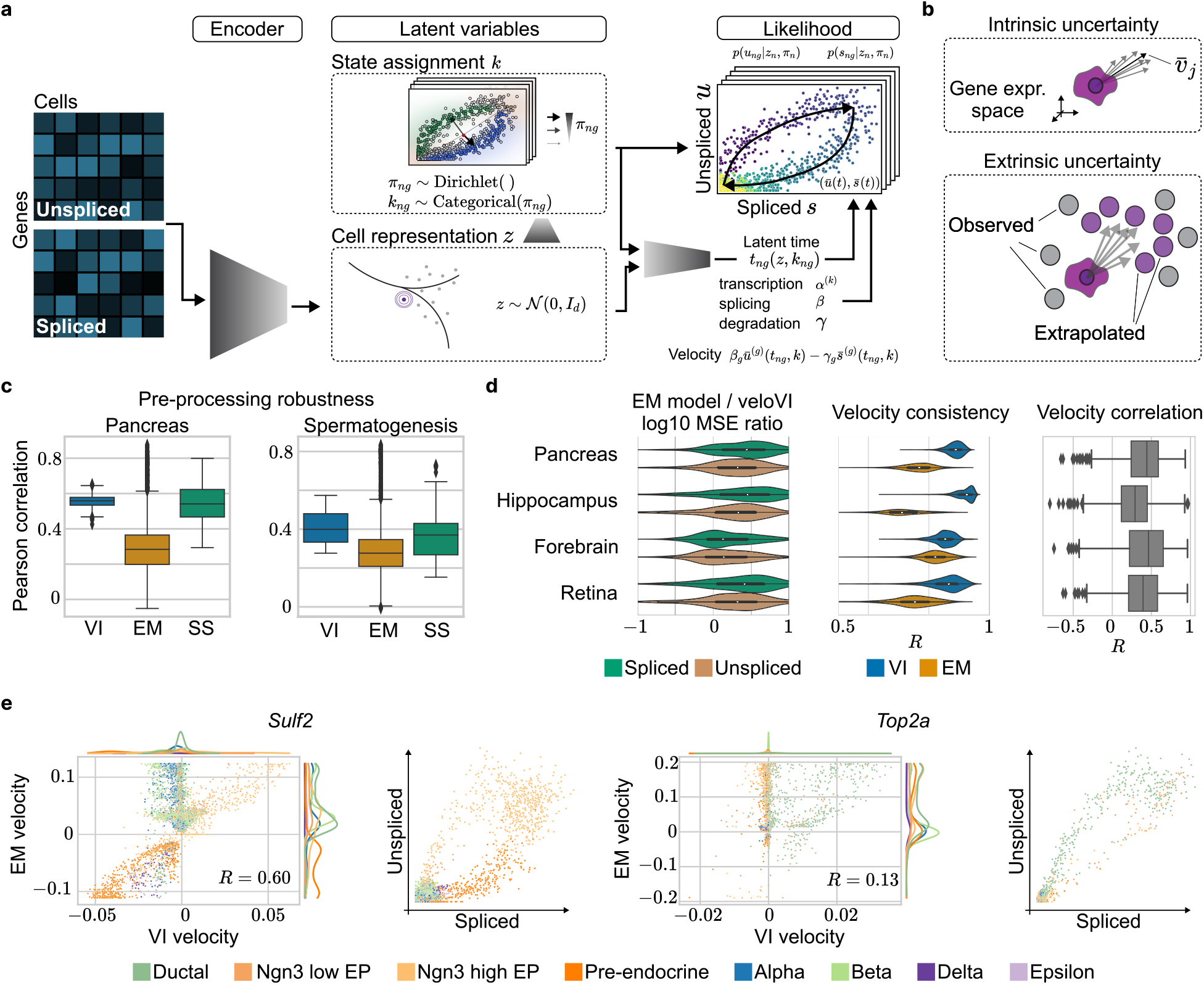
Overview of the veloVI model and benchmarks. **a**. veloVI encodes the unspliced and spliced abundances into the cell representation through a neural network. This cell representation is used to further encode the transcriptional state assignment of each cell/gene. Using the cell representation as input, the decoder neural network outputs a cell-gene-state specific latent time. The likelihood is a function of the latent time, kinetic rates (rates of transcription (*α*), splicing (*β*), and degradation (*γ*)), and uncertainty over the state assignment. **b**. The variational inference formulation quantifies the intrinsic uncertainty of a velocity estimate by sampling from the posterior distribution. This notion is contrasted by its extrinsic counterpart which quantifies variation within a cell’s neighborhood defined by transcriptomic similarity. **c**. Comparison of velocity estimation when using different algorithms to quantify unspliced and spliced counts. Velocity correlation of veloVI (VI), the *EM* (EM), and *steady-state model* (SS) are compared on the Pancreas^27^ and Spermatogenesis^28^ data. **d**. Comparison of veloVI and the *EM model* based on mean squared error (MSE) (left), velocity consistency (middle), and velocity correlation (right) on datasets of pancreas endocrinogenesis^27^, hippocampus^8^, forebrain^8^, and retina^29^. **e**. Velocity comparison on the level of individual genes in the Pancreas dataset (*Sulf2, Top2a*). For each gene, the velocity of the *EM model* is plotted against veloVI (left), and the gene phase portrait is given (right). Each observation is colored by its cell type as defined in previous work^27^.

As a Bayesian deep generative model, veloVI can output a posterior distribution over velocities at the cell-gene level. This distribution can be used to quantify an *intrinsic* uncertainty over first-order directions a cell can take in the gene space. In downstream analyses, velocity is often used to construct a cell-cell transition matrix that reweights the edges of a standard nearest-neighbors graph according to the similarity of the first-order displacement of a cell and its neighborhood^8,11^. By piping posterior velocity samples through this computation, we also quantify an *extrinsic* uncertainty, which reflects both the intrinsic uncertainty and the variability amongst the cell’s neighbors in gene space (Figure 1b; Methods). By contrast, the *EM model* and *steady-state model* do not carry any explicit notion of uncertainty. Indeed, both previous models only allow evaluating an uncertainty post-hoc based on quantifying velocity variation over a cell’s neighbors^9^. Finally, a point estimate of the velocity averaged over samples for a cell allows veloVI’s output to be used directly in scVelo’s downstream visualization and graph construction functionalities as well as other packages building upon scVelo^9,31^.

### veloVI improves data fit over the *EM model* and is stable

We performed a multifaceted analysis to evaluate veloVI’s ability to robustly fit transcriptional dynamics across a range of simulated and real datasets, comparing to both the *EM model* and the *steady-state* formulation of RNA velocity as implemented in the scVelo package^11^.

We first assessed each model’s ability to recover kinetic parameters in simulated data (Methods). With an increasing number of observations, we found that veloVI outperformed the *EM model* and was better than the steady-state model in recovering the simulated ratio of degradation and splicing rate for each gene (Supplementary Fig. 2a). We also benchmarked the runtime of veloVI and *EM model*. For this comparison, we ran both models on subsamples of a mouse retina dataset^32^ containing approximately 114,000 cells. Across multiple subsamples, inference was significantly faster using veloVI compared to the *EM model* (Supplementary Fig. 2b). Specifically, considering 20,000 cells, veloVI achieved a 5-fold speed-up.

We then evaluated the stability of velocity estimates on real datasets processed with 12 different flavors of RNA abundance quantification algorithms^8,32–36^, based on previous work that highlighted general inconsistencies in velocity estimation^37^ (Methods). To do so, we measured the correlation of velocity of each gene between pairs of quantification flavors on five benchmarking examples, namely pancreas endocrinogenesis at embryonic day 15.5^27^ as well as datasets of spermatogenesis^28^, mouse developing dentate gyrus^38^, the prefrontal cortex of a mouse^39^, and 21-22 months old mouse brain^40^. Averaging these correlations across all quantification algorithms, veloVI scored both a higher mean correlation and lower variance compared to the *EM model*. Compared to the much simpler *steady-state* model, veloVI tended to have a similar mean correlation, but with lower variance (Figure 1c, Supplementary Fig. 3-8).

To assess how well the inferred dynamics reflect the observed data, we computed the mean squared error (MSE) of the fit for the unspliced and spliced abundances and compared the MSE to that of the *EM model* on a selection of datasets (Supplementary Table 1). For each dataset, we computed the ratio of the MSE for veloVI and the *EM model* at the level of a gene. VeloVI had better performance for a majority of the genes in each dataset (Figure 1d). We then compared the local consistency of the velocity vector fields generated by each model. This consistency measure quantifies the extent to which the velocities of cells with similar transcriptomic profiles (i.e., nearest neighbors) agree. Across all datasets, veloVI had higher velocity consistency among cells (Figure 1d). We attribute this increase to the explicit low-dimensional modeling in veloVI that shares statistical strength across all cells and genes.

Despite sharing many model assumptions, the velocities estimated for a gene with veloVI were partially correlated on average with their *EM* counterpart (Figure 1d). To highlight the differences in velocity estimation at the level of individual genes, we examined *Sulf2*, a marker of endocrine progenitor cells, and *Top2a*, a cell cycle marker, in the pancreas dataset (Figure 1e). For both of these genes, the *EM model* predicted a wide range of velocities for cells that had near-zero unspliced and spliced abundances. For example, terminal Beta cells had substantially positive velocity under the *EM model* for *Sulf2* despite being located at the bottom-left of the phase portrait (defined as the scatter plot of unspliced versus spliced abundance of a gene) and with known development occurring later than endocrine progenitors and pre-endocrine cells. In the case of veloVI, Beta cells had nearly zero velocity, reflecting their belonging to the putative repression steady state for this gene. We attribute this result to veloVI*’s* velocity directly marginalizing over the latent cell representations, which explicitly incorporates the probability that a cell belongs to induction, repression, or their respective steady states (Methods). We observed similar results for *Top2a*, in which cell types without a strong cell-cycle signature and near-zero unspliced/spliced abundance had positive velocity in the *EM model*, but near-zero velocity using veloVI.

### veloVI enables interpretable velocity analysis

We then investigated how the uncertainty in the velocity estimates of veloVI could be used to scrutinize its output, both at the level of cells (which might be incorrectly modeled) and at the level of individual genes (which might be inconsistent with the aggregated, cell-level output). We used this uncertainty to (1) measure the variability in the phenotypic directionality suggested by the velocity vector in each cell (here, intrinsic uncertainty), and (2) quantify the variability of predicted future cell states under the velocity-induced cell-cell transition matrix (here, extrinsic uncertainty; Figure 1b; Methods).

We applied these uncertainty metrics to the pancreas dataset (Figure 2a). We observed that the intrinsic uncertainty was elevated in Ductal and Ngn3-low endocrine progenitor populations, while the extrinsic uncertainty highlighted these same populations in addition to terminal Alpha and Beta cells. These results demonstrate that lower intrinsic uncertainty does not necessarily preclude higher extrinsic uncertainty. While the former relies on estimating the velocity vector (which is cell-intrinsic), most velocity pipelines also account for other cells in the dataset, which presumably represent the potential past and future states of the cell, to determine cell transitions (Fig. 1b). In the case of Alpha and Beta cells, these cells represent terminal populations in the pancreas dataset, which may explain the high extrinsic uncertainty as there are no observed successor states. Conversely, in the case of transient cell populations, such as Ngn3-high endocrine progenitors and pre-endocrine cells, both metrics assign a low uncertainty. We attribute the low intrinsic uncertainty of these cells to the fact that their dynamics agree well with the underlying model assumptions (Supplementary Fig. 9). The addition of low extrinsic uncertainty further suggests that these cell types have clear successor populations in this dataset (Fig. 2a).

**Fig. 2.**
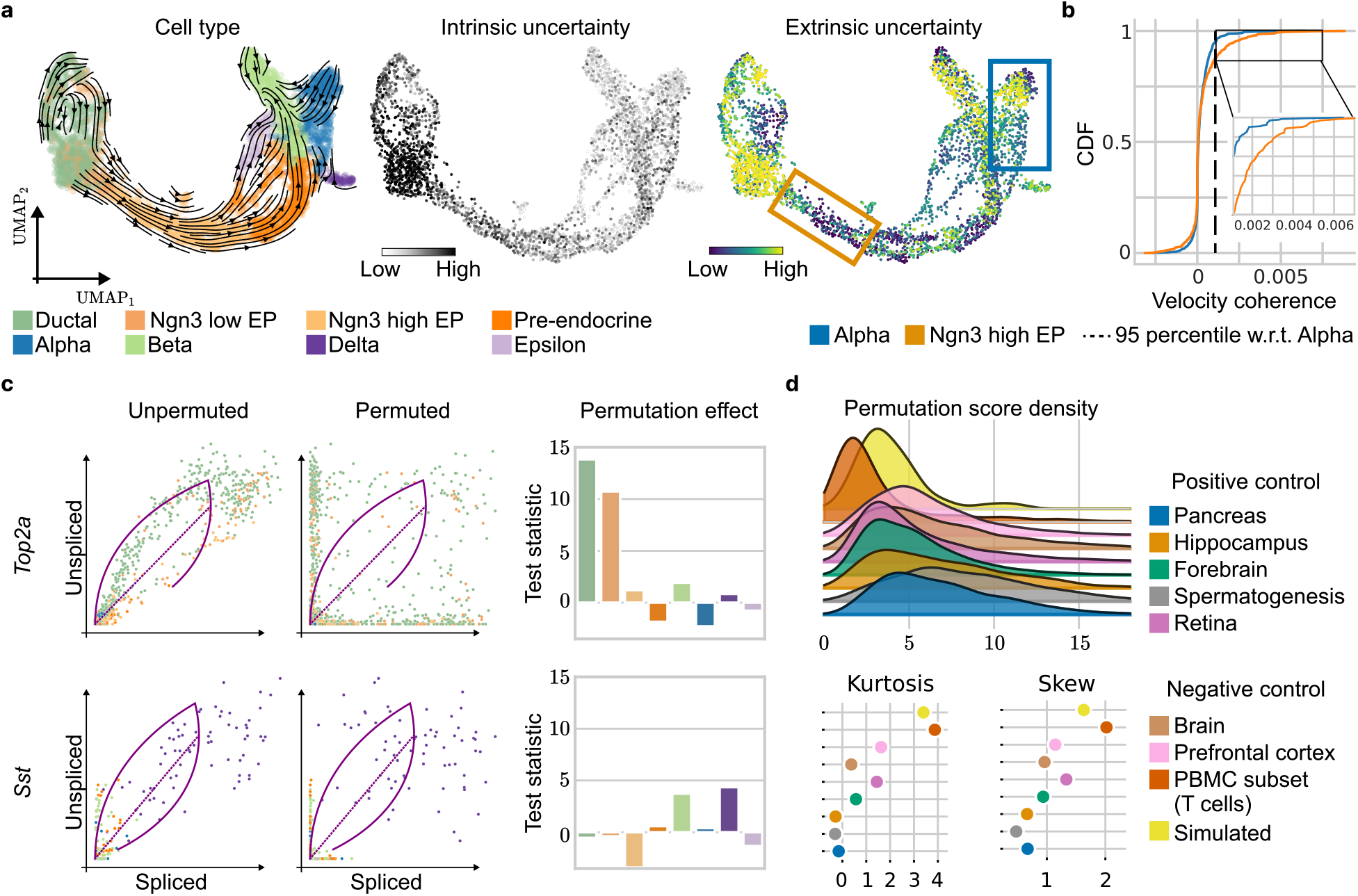
Uncertainty and permutation score analysis. **a**. From left to right: Velocity stream, intrinsic and extrinsic uncertainty of the Pancreas dataset estimated by veloVI. The left UMAP embedding is colored by cell type according to original cluster annotations^11,27^. Uncertainty is defined as the variance of the cosine similarity between samples of the velocity vector (intrinsic) and future cell states (extrinsic) and their respective mean (Methods). **b**. The corresponding cumulative distribution functions (CDFs) of the gene velocity coherence score is shown for Alpha and Ngn3-high EP cells. The velocity coherence, defined for one cell and gene as the product of the velocity and the expected displacement of that cell/gene, is averaged within cell types. **c**. The effect on the error between inferred dynamics and data when permuting unspliced and spliced abundance in the case of *Top2a* (top) and *Sst* (bottom). Coloring of cell types is according to panel a. **d**. Permutation score densities of datasets of the pancreas^27^, hippocampus^8^, forebrain^8^, spermatogenesis^28^, retina^29^, brain^40^, prefrontal cortex^39^, PBMC^41^, and simulated data^15^ (top). Kurtosis (left), and skew (right) for each dataset (bottom).

Finally, we ask whether we can use veloVI’s uncertainty to address the common behavior of unexpected “backflow” in two-dimensional velocity visualizations: When projecting the average veloVI velocity onto a UMAP^42^ plot (using procedures from ref.^11^), we observed an incorrect “backflow” of directionality in the Alpha and Beta cells, which showed transitions toward their known progenitors. While these terminal populations have high extrinsic uncertainty according to veloVI, it remains difficult to explain which genes cause the inconsistency. In the case of scVelo, it has been proposed to use the likelihood of a gene as a proxy, however, the likelihood has no direct connection to cell-cell transition-based analyses.

To this end, we sought to score genes in each cell according to how well their velocity agrees with the predicted future cell state that is derived via the velocity-induced transition matrix (i.e., incorporating velocity information from all genes as well as gene expression in neighboring cells; Methods). We reasoned that this score, which we call *velocity coherence*, could help gain insight into why a particular directionality might manifest. A positive score of a gene indicates the velocity value of that gene (i.e., the time derivative of its spliced mRNA) agrees with its expression in the inferred future cell state (same direction) and likewise, a negative score indicates disagreement (Figure 2b, Supplementary Fig. 10a).

In the Alpha cells, for example, there are both positively and negatively scoring genes. Genes with a negative score, like *Gcg* and *Sphkap*, were fit correctly by veloVI (Alpha cells after Pre-endocrine cells in time along the inferred trajectory on the phase portrait), but disagree with the predicted future cell state, suggesting that other genes are outweighing these genes in the transition matrix computation (Supplementary Fig. 10b). Indeed, genes like *Rnf130, Etv1*, and *Grb10*, which had a positive score that agrees with the backflow, appeared to have been fit incorrectly (Alpha cells precede Pre-endocrine cells along the inferred trajectory on the phase portrait) (Supplementary Fig. 10c). The incorrect fits can putatively be explained by violated model assumptions such as a transcriptional burst in Alpha cells (*Rnf130*), ambiguous phase portraits (*Etv1*), and multi-kinetics (*Grb10*).

Conversely, the dynamics in Ngn3-high cells are correctly visualized in the UMAP representation (Figure 2a). We attribute this result to the presence of many genes agreeing with both the model assumptions and the predicted future state of a cell (Supplementary Fig. 10d). Compared to the 95% percentile of the coherence score in Alpha cells, more than twice as many genes ranked above this threshold in the Ngn3-high cluster (135 vs. 54). However, even in this case, we found that many genes were fit with incorrect dynamics for this cell type (Supplementary Fig. 10e).

Taken together, these results suggest that the visualization of dynamics on a two-dimensional embedding with previously described procedures is explained by small subsets of genes. Thus, caution is warranted when analyzing projections of velocity estimates onto a two-dimensional embedding of the data. We urge users to investigate the dynamics at the level of individual genes to identify which genes meet the model assumptions. Putative candidates are given by our proposed velocity coherence score. Additionally, to identify genes viable for RNA velocity analysis due to the presence of transient cell populations, we propose a novel score outlined next.

### veloVI distinguishes genes and datasets with insufficiently observed or steady-state dynamics via permutation score densities

In datasets with non-differentiating, hierarchically-related cell types, spurious cell state transitions may manifest when applying RNA velocity^14,15^. Indeed, the underlying transcriptional likelihood model cannot readily distinguish between the case of a transient population and that of multiple steady-state populations. Therefore, we devised a procedure to use a trained veloVI model to identify genes with phase portraits that are consistent with a developmental process versus ones that are consistent with steady-state dynamics or are confounded by noise. For each gene and cell type, we first permuted the unspliced and spliced abundances of cells. We then inputted these permuted cells into the trained veloVI model and recorded the absolute error in the fit at the level of a gene and a cell type. Notably, veloVI can predict unspliced and spliced abundances for held-out cells, so the computation required for this procedure is trivial. To quantify the effect of the permutation on the model fit, we compared, for every cell type and for each gene, the absolute errors of permuted and unpermuted data with the t-test statistic (Methods). We hypothesized that the t-test statistic would be elevated in transient populations with strong time dependence, and, conversely, diminished in steady-state populations.

In the Pancreas dataset, the permutation strongly affects the Ductal and Ngn3 low EP cells for the cell-cycle gene *Top2a*. Indeed, these cell types trace fully-observed induction and repression states for *Top2a*. In the case of the Delta-specific gene *Sst*, where no such transient connection is observed, for example from Ductal to Pre-endocrine to Delta cells, no single cell type is strongly affected when permuting (Figure 2c). Consequently, even though *Sst* is essential for the identity of Delta cells, the gene does not display continuous dynamics from Ductal progenitor cells and, thus, does not include the necessary information to be analyzed with RNA velocity.

We then applied this procedure to a variety of datasets. In one set of tests, we used datasets describing cellular development. These datasets serve as partial positive controls since we expect directed dynamical processes, as modeled by RNA velocity, to take place in at least a subset of cells in the dataset. As negative controls, we used simulated data of bursty kinetics^15^ with no overall differentiation of cell state, and datasets containing multiple cell types that are in steady-state. To summarize the permutation for a gene, we used the maximum permutation effect statistic across cell types (permutation score). Two clusters of datasets emerged when characterizing the per-gene permutation score distribution (Figure 2d). One cluster, with a fatter right tail (quantified by skewness and kurtosis), contained positive control datasets like the Pancreas and Spermatogenesis. Despite having relatively many genes sensitive to permutation, the datasets of this cluster also contained many genes that were not sensitive, suggesting that there are likely many non-dynamical genes used for downstream analysis with RNA velocity. The other cluster, with less density in the right tail, contained negative controls like the PBMC, null data simulation, and the prefrontal cortex.

Between these two clusters of datasets, we also found a few ambiguous datasets, like the mouse retina (positive control) and brain (negative control), which suggests that there exist some cell subsets within these datasets that are affected by the permutation and hence, possibly reflect a directed dynamical process that is appropriate for modeling with RNA velocity. However, upon closer inspection of the brain dataset, we identified mature neurons as responsible for skewing the permutation score density (Supplementary Fig. 11a). The cluster of mature neurons was singled out as it attributes for about one-third of the highest permutation scores (Supplementary Fig. 11b). For the genes with the highest permutation score, these neuronal cells exhibit a bimodal distribution in which one mode has low unspliced and spliced abundance while the other has respectively higher abundances (Supplementary Fig. 11c). Thus, we attribute this skewing to coarse labeling of this population (Supplementary Fig. 11d). When excluding mature neurons from this analysis, the distribution shifted and its key characteristics moved towards the cluster formed by the negative control cases (Supplementary Fig. 11e).

In the accompanying code to this manuscript, we provide these permutation score densities as a resource for users of RNA velocity, which will enable the collection of datasets we analyzed here to serve as references for the distribution of these scores, and thus as a systematic approach to measure the overall transient dynamics of a dataset.

### veloVI is an extensible framework for modeling transcriptional dynamics

The transcriptional model assumptions at the level of one gene (e.g., constant rates that impose a specific structure of phase portraits) can be shown to be violated in many cases. For example, in the case of transcriptional bursts in which the transcription rate increases with time^18^ or multiple kinetics within a single gene^14^, the assumption of constant kinetic rates is violated. Thus, there remains a need for modeling frameworks that are extensible and support varied and more nuanced dynamical assumptions. While veloVI makes many of the same assumptions as in the *EM model*, it leverages black-box computational and statistical techniques that allow its generative model to be altered to include new assumptions without needing to extensively rewrite inference recipes or generally sacrifice scalability.

To explore veloVI as a general modeling framework, we adapted it to use gene-specific, time-dependent transcription rates. Under this extension, transcription rates are free to monotonically increase or decrease with respect to time^14^, thus allowing for modeling the acceleration of RNA abundance, which can impact the curvature of the model fit (Methods and Supplementary Fig. 12a). To infer these additional parameters, only the likelihood function of veloVI needed to be adapted. Applying this modified version of veloVI to the pancreas, dentate gyrus, and forebrain datasets, we observed improved fit for the majority of genes (Supplementary Fig. 12b). In the case of the pancreas dataset, the added flexibility allowed veloVI to better fit genes that appear more linear in their phase portraits, for example, as it can reduce the curvature of the fitted dynamics (Supplementary Fig. 12c).

In the case of *Smarca1*, the model using a constant transcription rate inferred a downregulation (repression) of Alpha cells differentiating into their progenitor populations of Pre-endocrine cell and Ductal cells (Supplementary Fig. 12c). Contrastingly, using a time-dependent transcription rate, the upregulation of Ductal to Pre-endocrine to Alpha cells is inferred by the generalized model. Similar observations apply to *Atad2* and *Cdkn1a*. While the constant transcription rate model inferred the correct regulation type for *Ppp1r1a*, its generalized counterpart captures the underlying dynamics more accurately (Supplementary Fig. 12d). Overall, for most genes, we observed a decreasing transcription rate over time (Supplementary Fig. 12e, f).

Altogether, this exemplary model extension demonstrates the flexibility of veloVI’s modeling approach. The flexibility allows to quickly prototype extensions and infer additional parameters within a single, consistent framework. We, thus, expect future models to benefit from such flexibility.

## Discussion

Here, we reformulated the estimation of RNA velocity in a variational inference framework with veloVI. This new method compares favorably to previously proposed methods^8,11^ and adds actionable metrics into downstream data analyses at the cell level via uncertainty quantification and at the level of a gene and dataset with the permutation score. In Supplementary Notes 1 and 2, we provide case studies outlining how veloVI can be used in practice on peripheral blood mononuclear cells (negative control) and mouse developing dentate gyrus (partial positive control). We believe that veloVI will facilitate more systematic analyses with RNA velocity and help reduce the strong reliance on prior knowledge to guide whether results are sensible. As an example, our permutation score could be used to filter genes that are considered for further analysis. We also note that related work has very recently incorporated deep learning with RNA velocity, and we review these methods and compare them to veloVI in Supplementary Note 3.

We view this formulation of modeling transcriptional dynamics with probabilistic models and deep learning as a step towards a more rigorous pipeline that faithfully captures the biophysical phenomenon of RNA metabolism. In this work, we relied on previously described data processing approaches that smooth unspliced/spliced abundances across nearest neighbors before velocity estimation. We also borrowed many assumptions from the *EM model*, including, for example, the lack of explicit support for multiple diverging lineages that would result in genes reflecting a superposition of dynamical signals. However, we built veloVI in an extensible way using the scvi-tools framework^43^. As a proof of concept, we demonstrated that veloVI could be easily extended to use time-dependent transcription rates, which improved model fit for many genes. We anticipate that the veloVI framework will be further adapted to overcome other computational challenges including estimating velocity while accounting for batch effects, using multi-modal technologies with measurements that span biology’s central dogma^44,45^, and directly modeling the unspliced and spliced RNA counts with count-based likelihoods.

A philosophical challenge with RNA velocity relates to the notion that models should use bottom-up mechanistic approaches while also being general enough to be applied across a variety of biological systems, each with their own caveats and unique dynamics. In this work, we use a low-dimensional representation of a cell’s phenotypic state to capture multiple biological processes (e.g., differentiation and cell cycle). More complex models likely need prior information, such as known experimental timepoints or cell type lineages to solve issues of statistical identifiability that arise in these more general modeling scenarios. However, incorporating such priors can contradict the usage of RNA velocity as a de novo discovery tool for the trajectory inference task. Despite all these outlined challenges, we envision that veloVI will facilitate applications of RNA velocity via uncertainty-aware analysis as well as easier model prototyping, benefiting both users and method developers.

## Methods

### veloVI model

For a full specification of the veloVI model, as well as how its outputs are used to compute uncertainties, please refer to the Supplementary Methods. Briefly, veloVI models the conditional distribution of unspliced and spliced RNA abundances for each gene in each cell as a mixture of Gaussian distributions with the mean of a mixture component defined by the transcription (*α*), splicing (*β*), and degradation (*γ*) rate parameters, latent time (*t*), and latent transcriptional state (*k*). This mean is explicitly defined by solving the system of ordinary differential equations used in the *EM model*:

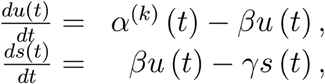

The latent transcriptional state distribution (*π*), which describes the probability of the induction, repression, induction steady-state, and repression steady-state, defines the mixture weights. Additionally, the latent time is a function of a low-dimensional (here, 10 dimensions) latent variable called the cell representation (*z*), where the function is defined as a fully-connected neural network called the decoder.

Inference consists of learning the posterior parameters over the latent variables, *z* and *π*, as well as estimating the global rate parameters and neural network parameters. This is done end-to-end in a variational inference framework in which the posterior parameters are parameterized using a series of encoder neural networks. veloVI is implemented using scvi-tools^43^, which enables GPU hardware acceleration during inference, as well as a user-friendly interface. Following inference, velocity is computed as a functional of the variational posterior following the ordinary differential equation:

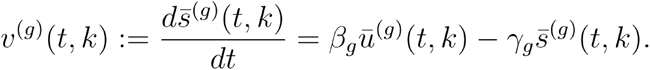

Given that there are posterior distributions over the latent variables, veloVI can output posterior samples of velocity at the level of a cell-gene pair. We used samples of this distribution for the uncertainty experiments. As a point estimate for velocity, we use the mean over the samples. This is used for any comparison against the *EM* or *steady-state models*.

### Data preprocessing

All datasets were pre-processed following the same steps. Genes with fewer than 20 unspliced or spliced counts are removed. Transcriptomic counts of each cell are normalized by their median, pre-filtered library size and the 2000 most highly variable genes selected based on dispersion. The aforementioned steps are performed using scVelo’s^11^ `filter_and_normalize` function.

Following gene filtering and count normalization, the first 30 principal components are calculated and a nearest neighbor graph with *k* = 30 neighbors is constructed. In a final step, counts are smoothed by the mean expression across their neighbors in order to compute final RNA abundances. These steps are performed by scVelo’s `moments` function.

To estimate RNA velocity, the preprocessed unspliced and spliced abundances are (gene-wise) min-max scaled to the unit interval. Following, the *steady-state model* is applied to the entire dataset. Genes for which the estimated steady-state ratio and *R*^2^ statistic are positive are considered for further analysis. If not stated otherwise, this subset of genes is used for parameter inference of veloVI and the *EM model*.

### Benchmarking against EM and steady-state models

VeloVI was benchmarked against the *EM* and *steady-state model* by first comparing the accuracy of inferred parameters on simulated data. For each number of observations (1000, 2000, 3000, 4000, 5000), we simulated ten datasets of unspliced and spliced counts with 1000 kinetic parameters following a log-normal distribution. Latent time is Poisson distributed with a maximum of 20 hours with the switch from induction to transcription taking place after two to ten hours. The simulations were performed using the `simulation` function as implemented in scVelo^11^ with `noise_level=0.8`.

To compare the runtimes of veloVI and *EM model* were run on random subsets a mouse retina dataset^13^ containing 1000, 3000, 5000, 7500, 10000, 15000, 20000 cells. The *EM model* was run on an Intel(R) Core(TM) i9-10900K CPU @ 3.70GHz CPU using 8 cores. VeloVI was run on an Nvidia RTX3090 GPU.

In the case of real-world data, for each gene, the mean squared error (MSE) for unspliced and spliced counts (*u, s*) using the *VI* and *EM model* are compared. We compute for each model and modality the MSE given by

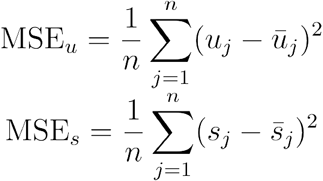

where 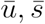 denote the estimated quantities. In the case of the *EM model*, this is directly a function of the rates, time, and transcriptional state, and in the case of veloVI this is the posterior predictive mean (Supplementary Methods).

In addition to the MSE, the model-specific velocity consistency^11^ can be compared. The velocity consistency quantifies the mean correlation of the velocity *υ*(*x*_*j*_) of a reference cell *x*_*j*_ with the velocities of its neighbors *𝒩*_*k*_(*x*_*j*_) in a k-nearest neighbor graph.

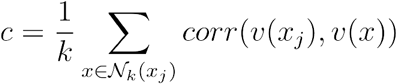

To calculate the consistency, we rely on scVelo’s `velocity_confidence` function.

Velocity estimates *υ*^(VI)^ and *υ*^(EM)^ of the *VI* and *EM model*, respectively, are compared through Pearson correlation given by

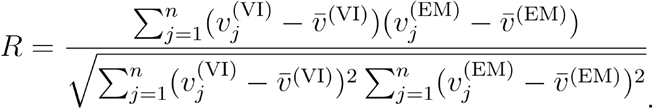

### Stability analysis across quantification algorithms

To assess the robustness of estimation using different means of quantifying unspliced and spliced reads, we relied on previously preprocessed and published data^37^. The collection contains outputs of variants of the alevin^33^, kallist/bustools^33,46,47^, velocyto^8^, dropEST^36^, and starsolo^48^ pipelines. For details of how the data was generated, we refer to the original work^37^.

To compare estimation across quantification algorithms, we first defined a reference set of genes for which to calculate RNA velocity. The set of reference genes was defined as the set of genes kept by preprocessing the data of one quantification method. In the case of the dentate gyrus data, starsolo was chosen for the quantification method, for all others velocyto. Data was pre-processed according to our described preprocessing pipeline. Counts from all other quantification approaches the same pre-processing steps were followed except for gene filtering. To prevent the reference genes from being filtered out, they are passed to the `filter_and_normalize` function via the argument `retain_genes`.

Velocities were estimated for the *steady-state, EM*, and veloVI. The velocities of the first two models were quantified using the function `velocity` with `mode=“deterministic”` and `mode=“dynamics”`, respectively, implemented in scVelo^11^. For veloVI, model parameters were inferred using default parameters and mean velocities estimated from 25 samples drawn from the posterior.

To compare estimates across quantification algorithms, for each model, cell, and pair of quantification algorithms, the Pearson correlation between the corresponding velocity estimates *υ*^(1)^, *υ*^(2)^, respectively,

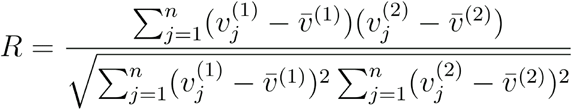

is calculated. The correlation scores can be aggregated to assess robustness for each model.

### Analysis with uncertainty quantification and velocity coherence

VeloVI allows estimating an intrinsic and extrinsic uncertainty (Supplementary Methods). The intrinsic uncertainty is quantified by first sampling the velocity vector 100 times from the posterior distribution. Next, for each cell, the cosine similarity between each sample and their mean is calculated. The variance of this distribution gives the intrinsic uncertainty.

The extrinsic uncertainty for each cell is defined similarly to its intrinsic counterpart. Instead of generating velocity samples from the posterior distribution, we compare the expected future states of a given cell. Future states of a cell are defined through the transition matrix *T*(*υ, s*) as described for the *EM model* in our previous work^14^. Given one sample of velocity, we compute *T*(*υ, s*)*S*, where *T*(*υ*) is the cell-cell transition matrix and *S* is the cell by genes matrix of spliced RNA abundances. The predicted future cell states here are a function of one sample of velocity, and uncertainty is then quantified using the same procedure as the intrinsic uncertainty with these cell state vectors.

Extrapolated future states *Tc*_*j*_ of a cell can also be used to evaluate if inferred velocities are coherent. The velocity *υ*_*j*_ of a given cell *c*_*j*_ is coherent if it points in the same direction as the empirical displacement *δ*_*j*_ = *Tc*_*j*_ − *c*_*j*_. Directionality is compared by calculating the Hadamard product *δ*_*j*_ *○ υ*_*j*_. In case both vectors point in the same direction for a given cell, the resulting entry will be positive and negative otherwise. To aggregate the score we report its mean per gene and cell type.

### Permutation scoring

To quantify how robust the inferred dynamics are with respect to random permutations in the input data, we define a gene- and cell-type-specific permutation score. For this analysis, we consider all highly variable genes and do not filter our genes based on estimates of the steady-state model. To calculate the score, for each gene and cell type, the unspliced and spliced abundance are independently permuted: Given the unspliced (respective spliced) abundance *u* (resp., *s*) of one gene, we permute its indices of the observations within each cell type with a permutation *π*_*u*_ (resp., *π*_*s*_). The permutation defines the permuted vector *u*^(*p*)^

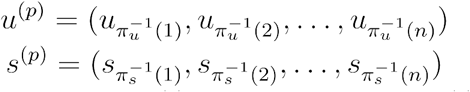

We then take the absolute error between *u*^(*p*)^ and the model fit of *u*^(*p*)^ (i.e., the posterior predictive mean, Supplementary Methods) and similarly for *s*^(*p*)^, summing the unspliced and spliced errors per newly permuted cell. Note that because veloVI can handle held-out data, computing the model fit of permuted data does not require any additional training. The mean absolute error between permuted observations and inferred dynamics is denoted by *μ*_*p*_. Repeating this error computation for unpermuted data, we denote the mean absolute error for unpermuted data as *μ*_0_. The permutation does not affect results if the two statistics are equal. We reason that this scenario would occur for a completely steady-state population.

To test if the mean absolute errors of the two samples are equal, we define the permutation score as the independent, cell-type specific t-test statistic

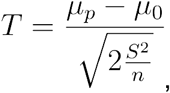

with number of cells *n*, and pooled variance *S*^2^ of the absolute errors. In order to limit the effect of dataset size, we consider the maximum sample size *n* = 200 of observations. The permutation score is aggregated on a gene level by considering the maximum test statistic across cell types. This aggregation allows comparing the permutation score across different datasets.

### Time-dependent transcription rates

Assuming a constant kinetic rate is one assumption of both the *steady-state* and *EM model*. To highlight the model extensibility of veloVI, the model can be fitted with a time-dependent transcription rate defined as

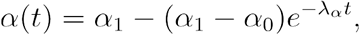

instead. Parameters *α*_0_, *α*_1_, *λ*_*α*_ are inferred by the model. The resulting equations for unspliced and spliced counts read

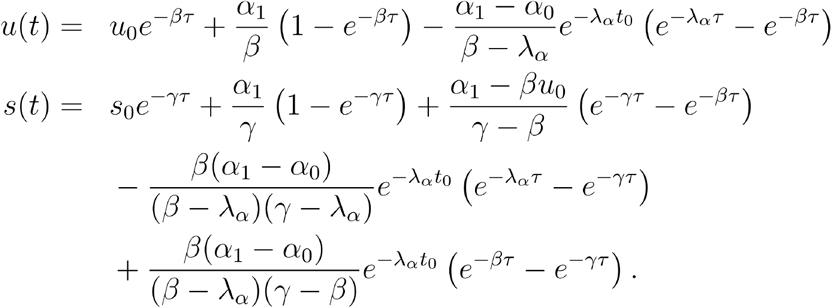

## Code and data availability

veloVI is currently implemented in a standalone package at https://github.com/YosefLab/velovi. The authors will move the core veloVI model into scvi-tools (https://scvi-tools.org/) during reviews in addition to providing a wrapper in the scVelo package (https://scvelo.org/). Code to reproduce the results in the manuscript can be found at: https://github.com/YosefLab/velovi_reproducibility.

Processed data, including spliced and unspliced count abundances, is available via scVelo, the Friedrich Miescher Institute for Biomedical Research (https://www.fmi.ch/groups/gbioinfo/RNAVeloQuant/RNAVeloQuant.html), and figshare (https://figshare.com/projects/veloVI_datasets/145476).

## Acknowledgments

We thank Romain Lopez and Matt Jones for feedback on the concepts and benchmarking of veloVI. We acknowledge members of the Streets, Theis, and Yosef laboratories for general feedback. A.S. is a Chan Zuckerberg Biohub investigator. A.G. and N.Y were supported by the Chan Zuckerberg Initiative Essential Open Source Software Cycle 4 grant (EOSS4-0000000121) for scvi-tools. M.L. acknowledges financial support from the Joachim Herz Stiftung via Add-on Fellowships for Interdisciplinary Life Science. F.J.T. acknowledges support by the BMBF (grant #01IS18036B and grant #01IS18053A) and by the Helmholtz Associations Initiative and Networking Fund through Helmholtz AI [grant #ZT-I-PF-5-01].

## Author contributions

A.G. and P.W. contributed equally. A.G., P.W., and M.L. conceptualized the study. A.G. conceptualized the statistical model with contributions from M.L. and P.W.. A.G. designed and implemented veloVI with contributions from P.W., J.H., and M.L. P.W. designed and implemented modeling extensions. D.K. designed and implemented model uncertainty analyses with contributions from A.G., P.W., and M.L.. A.G., P.W., and J.H. designed and implemented analysis methods with contributions from M.L.. A.S., F.J.T., and N.Y. supervised the work. A.G., P.W., M.L., F.J.T., and N.Y. wrote the manuscript.

