## Supplementary information for "Deep generative modeling of transcriptional dynamics for RNA velocity analysis in single cells"

**Adam Gayoso [1, *], Philipp Weiler [2, 3, *], Mohammad Lotfollahi [2], Dominik Klein [2], Justin Hong [1], Aaron Streets [1, 4, 5], Fabian J. Theis [2, 3, 6, #], and Nir Yosef [1, 7, #]**

[1] Center for Computational Biology, University of California, Berkeley, Berkeley, USA
[2] Institute of Computational Biology, Helmholtz Center Munich, Munich, Germany
[3] Department of Mathematics, Technical University of Munich, Munich, Germany
[4] Department of Bioengineering, University of California, Berkeley, Berkeley, USA
[5] Chan Zuckerberg Biohub, San Francisco, USA
[6] TUM School of Life Sciences Weihenstephan, Technical University of Munich, Munich, Germany
[7] Department of Electrical Engineering and Computer Sciences, University of California, Berkeley, Berkeley, USA

* These authors contributed equally.

**Supplementary Tables** **24**

### Supplementary Methods

#### Model specification

We begin with the formulation of the "dynamical" model of RNA velocity as presented by ref. [1]. We posit transcriptional states $k \in \{1, 2, 3, 4\}$, where $k = 1$ indicates induction, $k = 2$ indicates the induction steady state, $k = 3$ indicates repression, and $k = 4$ indicates the repression steady state.

Let $\alpha_{gk}$ be the gene-state-specific reaction rate of transcription. Let $\beta_g$ be the gene specific splicing rate constant and let $\gamma_g$ be the gene specific degradation rate constant. Each gene has a switching time $t_g^s$ when the system switches from induction phase to repression phase.

Given the solution to the ordinary differential equations in ref. [1], the unspliced transcript abundance at time $t_{ng}$ for cell $n$ and gene $g$ is defined as

$$\bar{u}^{(g)}(t_{ng}, k) := u_{gk}^0 e^{-\beta_g (t_{ng} - t_{gk}^0)} + \frac{\alpha_{gk}}{\beta_g} \left(1 - e^{-\beta_g (t_{ng} - t_{gk}^0)}\right), \tag{1}$$

where $t_{gk}^0$ is the initial time of the system in state $k$. The spliced transcript abundance is defined as

$$\begin{aligned} \bar{s}^{(g)}(t_{ng}, k) :=& s_{gk}^0 e^{-\gamma_g \tau} + \frac{\alpha_{gk}}{\gamma_g} \left(1 - e^{-\gamma_g (t_{ng} - t_{gk}^0)}\right) \\ &+ \frac{\alpha_{gk} - \beta_g u_{gk}^0}{\gamma_g - \beta_g} \left(e^{-\gamma_g (t_{ng} - t_{gk}^0)} - e^{-\beta_g (t_{ng} - t_{gk}^0)}\right). \end{aligned} \tag{2}$$

**Induction state** For the induction state, $k = 1$, we have $u_{g1}^0 = 0$, $s_{g1}^0 = 0$, $\alpha_{g1} > 0$ and $t_{g1}^0 = 0$. Thus, the unspliced transcript abundance can then be expressed as

$$\bar{u}^{(g)}(t_{ng}, k = 1) := \frac{\alpha_{g1}}{\beta_g} \left(1 - e^{-\beta_g t_{ng}}\right). \tag{3}$$

Likewise, the spliced transcript abundance can be simplified to

$$\bar{s}^{(g)}(t_{ng}, k = 1) := \frac{\alpha_{g1}}{\gamma_g} \left(1 - e^{-\gamma_g t_{ng}}\right) + \frac{\alpha_{g1}}{\gamma_g - \beta_g} \left(e^{-\gamma_g t_{ng}} - e^{-\beta_g t_{ng}}\right). \tag{4}$$

**Induction steady state** For the induction steady state, $k = 2$, the unspliced and spliced transcript abundances are defined as limits of the system:

$$\bar{u}^{(g)}(t_{ng}, k = 1) := \lim_{t_{ng} \to \infty} \bar{u}^{(g)}(t_{ng}, k = 1) = \frac{\alpha_{g1}}{\beta_g} \tag{5}$$

$$\bar{s}^{(g)}(t_{ng}, k = 2) := \lim_{t_{ng} \to \infty} \bar{s}^{(g)}(t_{sg}, k = 1) = \frac{\alpha_{g1}}{\gamma_g}. \tag{6}$$

**Repression state** For the repression state, $k = 3$, we have $\alpha_{g3} = 0$ and $t_{g3}^0 = t_g^s$. Thus, the number of unspliced transcripts can then be expressed as

$$\bar{u}^{(g)}(t_{ng}, k = 3) := u_{g3}^0 e^{-\beta_g (t_{ng} - t_{g3}^0)}. \tag{7}$$

Likewise, the number of spliced transcripts can be simplified to

$$\bar{s}^{(g)}(t_{ng}, k = 3) := s_{g3}^0 e^{-\gamma_g (t_{ng} - t_{g3}^0)} - \frac{\beta_g u_{g3}^0}{\gamma_g - \beta_g} \left(e^{-\gamma_g \tau} - e^{-\beta_g (t_{ng} - t_{g3}^0)}\right). \tag{8}$$

The initial conditions, $u^0_{g3}$ and $s^0_{g3}$ are defined by the induction model at the switching time $t^s_g$, such that

$$u^0_{g3} = \bar{u}^{(g)}(t_{sg}, k = 2) \tag{9}$$

$$s^0_{g3} = \bar{s}^{(g)}(t_{sg}, k = 2). \tag{10}$$

**Repression steady state** For the repression steady state, the limit upon which $t_{ng} \to \infty$, there is no expression, so we have that

$$\bar{u}^{(g)}(t_{ng}, k = 4) := 0 \tag{11}$$

$$\bar{s}^{(g)}(t_{ng}, k = 4) := 0. \tag{12}$$

**Model assumptions** As in ref. [1], this model assumes that for one gene, at the initial time of the system, cells are first in induction phase in which both spliced and unspliced expression increases. Then cells potentially reach a steady state of this induction state. Next at some future time $t^s_g$ the system switches to repression state. Finally, the repression reaches a steady state in which there is no expression. Further assumptions are necessary to identify the dynamical model parameters [2]; thus, we assume that each gene is on the same time scale (precisely each gene has a maximum time of $t = 20$ as in [1]).

#### Generative process

We posit a generative process that takes into account the underlying dynamics of the system. Compared to Bergen et al. [1], the model here does not treat each gene independently; instead, the latent time and states for each (cell, gene) pair are tied together via a local low-dimensional latent variable.

**For each cell** we draw a low-dimensional (10 dimensions throughout this manuscript) latent variable

$$z_n \sim \text{Normal}(0, I_d) \tag{13}$$

that summarizes the latent state of each cell. Next, **for each gene** $g$ **in cell** $n$ we draw the distribution over the state assignments as well as the state assignment itself

$$\pi_{ng} \sim \text{Dirichlet}(0.25, 0.25, 0.25, 0.25) \tag{14}$$

$$k_{ng} \sim \text{Categorical}(\pi_{ng}) \tag{15}$$

If $k_{ng} = 1$ (induction), then the time is a function of $z_n$,

$$\rho^{(1)}_{ng} = [h_{\text{ind}}(z_n)]_g \tag{16}$$

$$t^{(1)}_{ng} = \rho^{(1)}_{ng} t^s_g \tag{17}$$

where $h_{\text{ind}} : \mathbb{R}^d \to (0, 1)^G$ is parameterized as a fully-connected neural network. Notably, this parameterization results in an induction-specific time that is constrained to be less than the switching time.

Else, if $k_{ng} = 3$ (repression),

$$\rho^{(3)}_{ng} = [h_{\text{rep}}(z_n)]_g \tag{18}$$

$$t^{(3)}_{ng} = (t_{\max} - t^s_g)\rho^{(3)}_{ng} + t^s_g \tag{19}$$

where $t_{\max} := 20$ is used to fix the time scale across genes and identify the rate parameters of the model. Similarly to the previously defined function, $h_{\text{rep}} : \mathbb{R}^d \to (0,1)^G$, and is also a neural network.

We also consider two potential steady states. If $k_{ng} = 2$ (induction steady state) or if $k_{ng} = 4$ (repression steady state), we consider the limit as time approaches $\infty$, which was described in the previous section.

Finally, the observed data are sampled from normal distributions as

$$u_{ng} \sim \text{Normal}(\bar{u}^{(g)}(t_{ng}^{(k_{ng})}, k_{ng}), (c_k \sigma_g^u)^2) \tag{20}$$

$$s_{ng} \sim \text{Normal}(\bar{s}^{(g)}(t_{ng}^{(k_{ng})}, k_{ng}), (c_k \sigma_g^s)^2) \tag{21}$$

For veloVI, we consider the observed data to be the nearest-neighbor smoothed expression data that is also used as input to scVelo as well as velocyto. In addition, we assume the data have been preprocessed such that for each gene, the smoothed spliced and unspliced abundances are independently min-max scaled into $[0, 1]$. However, we feel that the flexibility of this modeling framework will enable extensions that consider the discrete nature of unique molecular identifiers used in standard scRNA-seq assays. We include a state-dependent scaling factor on the variance. For all experiments in this manuscript we used $c_k = 1$ except for the repression steady state in which $c_4 = 0.1$. In the following, let $\theta$ be the set of parameters of the generative process ($\alpha, \beta, \gamma, t^s$, and neural network parameters).

#### Inference

We seek the following: (1) point estimates of the transcription rate, degradation and splicing rate constants, and the switching time point, (2) point estimates of the parameters of the neural networks, and (3), a posterior distribution over the latent variables, which in this case includes $z$ and $\pi$. Noting that the model evidence $p_\theta(u, s)$ cannot be computed in closed form, we use variational inference [3] to approximate the posterior distribution as well as accomplish the other tasks. Following inference, velocity can be calculated as a functional of the variational posterior distribution.

**Variational posterior** We posit the following factorization on the approximate posterior distribution

$$q_\phi(z, \pi \mid u, s) := \prod_n^N q_\phi(z_n \mid u_n, s_n) \prod_g^G q_\phi(\pi_{ng} \mid z_n), \tag{22}$$

in which dependencies are specified using neural networks with parameter set $\phi$.

For the likelihoods, we integrate over the choice of transcriptional state $k_{ng}$, such that the likelihoods for unspliced and spliced transcript abundances,

$$p_\theta(u_{ng} \mid z_n, \pi_n) = \sum_{k_{ng} \in \{1,2,3,4\}} \pi_{ngk_{ng}} \text{Normal}(\bar{u}^{(g)}(t_{ng}^{(k_{ng})}, k_{ng}), (c_k \sigma_g^u)^2) \tag{23}$$

$$p_\theta(s_{ng} \mid z_n, \pi_n) = \sum_{k_{ng} \in \{1,2,3,4\}} \pi_{ngk_{ng}} \text{Normal}(\bar{s}^{(g)}(t_{ng}^{(k_{ng})}, k_{ng}), (c_k \sigma_g^s)^2) \tag{24}$$

are mixtures of normal distributions.

**Objective** The objective that is minimized during inference is composed of two terms

$$\mathcal{L}_{\text{velo}}(\theta, \phi; u, s) = \mathcal{L}_{\text{elbo}}(\theta, \phi; u, s) + \lambda \mathcal{L}_{\text{switch}}(\theta; u, s), \tag{25}$$

where $\mathcal{L}_{\text{elbo}}$ is the negative evidence lower bound [3] of $\log p_\theta(u, s)$ and $\mathcal{L}_{\text{switch}}$ is an additional penalty that regularizes the location of the transcriptional switch in the phase portrait. In more detail,

$$\mathcal{L}_{\text{elbo}}(\theta, \phi; u, s) = \sum_n -\mathbb{E}_{q_\phi(z_n, \pi_n \mid u_n, s_n)} \left[\log p_\theta(u_n, s_n \mid z_n, \pi_n)\right] + \text{KL}\left(q_\phi(z_n \mid u_n, s_n) \parallel p(z)\right) + \mathbb{E}_{q_\phi(z_n \mid u_n, s_n)} \left[\sum_g \text{KL}\left(q_\phi(\pi_{ng} \mid z_n) \parallel p(\pi_{ng})\right)\right], \tag{26}$$

which can be estimated using minibatches of data. In particular, we use minibatches of 256 cells for inference. For the penalty term $\mathcal{L}_{\text{switch}}$, we start by only considering cells that are above the 99th percentile of unspliced abundance for each gene. Using these cells we compute the median unspliced and spliced abundance for each gene separately. Let $u^*$ and $s^*$ be the outcome of this procedure, then

$$\mathcal{L}_{\text{switch}}(\theta; u, s) = \sum_g \left(u^0_{g3} - u^*_g\right)^2 + \left(s^0_{g3} - s^*_g\right)^2, \tag{27}$$

where $u^0_{g3}$ and $s^0_{g3}$ were defined as the initial conditions of the repression phase at the switch time $t^s_g$.

**Initialization** We initialize $\alpha_{g1}$ to be equal to the median unspliced abundance for the cells above the 99th percentile for each gene. The other global parameters, including the splicing, degradation, and switch time are initialized to a constant value shared by all genes. All neural network initialization uses the default implementation in PyTorch.

**Optimization** To optimize $\mathcal{L}_{\text{velo}}$ we use stochastic gradients [4] along with the Adam optimizer with weight decay [5] as implemented in PyTorch [6]. For all experiments we use $\lambda = 0.2$ for scaling the regularization term in the loss. As a result of minibatching, veloVI's memory usage is constant throughout training. Unless otherwise specified, all neural networks are fully-connected feedforward networks that use standard activation functions like ReLU for hidden layers and softplus or exponential for parameterizing non-negative distributional parameters.

#### Downstream tasks

**Fitted abundance values** The fitted values (used for e.g., mean squared error benchmarks) for unspliced and spliced abundance are the posterior predictive mean:

$$\mathbb{E}_{p(u^*_n \mid u_n, s_n)}\left[u^*_n\right], \quad \mathbb{E}_{p(s^*_n \mid u_n, s_n)}\left[s^*_n\right],$$

with posterior predictive in the case of unspliced defined as

$$p(u^*_n \mid u_n, s_n) = \mathbb{E}_{q_\phi(z_n, \pi_n \mid u_n, s_n)}\left[p_\theta(u^*_n \mid z_n, \pi_n)\right],$$

which uses the variational posterior distribution as a plug-in estimator for the true (unknown) posterior distribution.

**State assignment** The state assignment for each gene and cell is the approximate posterior mean

$$\mathbb{E}_{q_\phi(z_n \mid u_n, s_n)}\left[\mathbb{E}_{q_\phi(\pi_{ng} \mid z_n)}[\pi_{ng}]\right].$$

**Gene-wise latent time** The latent time is computed for each gene and cell as

$$\mathbb{E}_{q_\phi(z_n \mid u_n, s_n)}\left[\mathbb{E}_{q_\phi(\pi_{ng} \mid z_n)}[t^{(k_{ng})}_{ng}]\right],$$

where the outer expectation with respect to $q_\phi(z_n \mid u_n, s_n)$ is estimated with Monte Carlo samples, while the inner expectation is computed analytically over the transcriptional states $k_{ng}$.

**RNA velocity** The velocity of a particular gene in a particular cell is similarly a functional of the variational posterior. Recall that the velocity is computed as

$$v^{(g)}(t^{(k)}, k) := \frac{d\bar{s}^{(g)}(t, k)}{dt}\Big|_{t^{(k)}} = \beta_g \bar{u}^{(g)}(t^{(k)}, k) - \gamma_g \bar{s}^{(g)}(t^{(k)}, k).$$

Thus, we can compute samples of a posterior predictive velocity distribution via the following process

1. Sample $z_n$ from $q_\phi(z_n \mid u_n, s_n)$.
2. Compute $\mathbb{E}_{q_\phi(\pi_{ng} \mid z_n)}[v^{(g)}(t_{ng}^{(k_{ng})}, k_{ng})]$ for each gene.

This provides samples from a distribution over the velocity for every gene, cell pair, which we then use in downstream tasks.

**Intrinsic uncertainty** Let $\bar{v}_n$ be the posterior predictive velocity mean from the procedure above. The intrinsic uncertainty is then computed as $\mathbb{V}\mathrm{ar}_{q_\phi(v_n \mid u_n, s_n)}[c(v_n, \bar{v}_n)]$ where $c$ denotes the cosine similarity. In effect, denote by $\{v_n^{(l)}\}_{l=1}^L$ the set of $L$ velocity vector samples of cell $n$ from the variational posterior. Then we have:

$$\hat{\sigma}_n^2 = \frac{1}{L-1} \sum_{l=1}^{L} \Big( \frac{v_n^{(l)} \cdot \bar{v}_n}{||v_n^{(l)}||\,||\bar{v}_n||} - \frac{1}{L} \sum_{j=1}^{L} \frac{v_n^{(j)} \cdot \bar{v}_n}{||v_n^{(j)}||\,||\bar{v}_n||} \Big)^2. \tag{28}$$

**Extrinsic uncertainty** Let $T(v_{1:N}, s_{1:N})$ be a function that maps the velocity vectors and spliced abundances of the entire dataset (with $N$ cells) to a cell-cell transition matrix computed as described in ref. [1]. Namely, this function compares the similarity of the displacement $\delta_{ij}$ of nearest neighbors $s_i$ and $s_j$ (defined using $s_{1:N}$) to the velocity of cell $i$, $v_i$, via the cosine similarity

$$\cos(\delta_{ij}, v_i) = \frac{\delta_{ij}^T v_i}{||\delta_{ij}||||v_i||} \tag{29}$$

as the basis for computing transition probabilities between pairs of cells.

Following the construction of $T(v_{1:N}, s_{1:N})$ for one sample of velocity, the predicted future cell state is computed by the matrix multiplication $T(v_{1:N}, s_{1:N})S$, where $S$ is the cells by genes matrix of spliced RNA abundances. These predicted future cell state vectors (over samples of velocity) then undergo the same variance computation procedure as described for the intrinisic uncertainty (namely, variance of the cosine similarity).

#### Time-dependent transcription rate

To showcase veloVI's model extensibility, we consider the system

$$\begin{aligned} \dot{u} &= \alpha^{(k)} - \beta u \\ \dot{s} &= \beta u - \gamma s, \end{aligned} \tag{30}$$

with

$$\alpha = \alpha(t) = \alpha_1 - (\alpha_1 - \alpha_0)e^{-\lambda_\alpha t}. \tag{31}$$

The system is solved by

$$
\begin{aligned}
u(t) = &\quad u_0 e^{-\beta\tau} + \frac{\alpha_1}{\beta}\left(1 - e^{-\beta\tau}\right) - \frac{\alpha_1 - \alpha_0}{\beta - \lambda_\alpha} e^{-\lambda_\alpha t_0}\left(e^{-\lambda_\alpha \tau} - e^{-\beta\tau}\right) \\
s(t) = &\quad s_0 e^{-\gamma\tau} + \frac{\alpha_1}{\gamma}\left(1 - e^{-\gamma\tau}\right) + \frac{\alpha_1 - \beta u_0}{\gamma - \beta}\left(e^{-\gamma\tau} - e^{-\beta\tau}\right) \\
&- \frac{\beta(\alpha_1 - \alpha_0)}{(\beta - \lambda_\alpha)(\gamma - \lambda_\alpha)} e^{-\lambda_\alpha t_0}\left(e^{-\lambda_\alpha \tau} - e^{-\gamma\tau}\right) \\
&+ \frac{\beta(\alpha_1 - \alpha_0)}{(\beta - \lambda_\alpha)(\gamma - \beta)} e^{-\lambda_\alpha t_0}\left(e^{-\beta\tau} - e^{-\gamma\tau}\right)
\end{aligned}
\tag{32}
$$

and the exact derivation given in Supplementary Note 4.

### Supplementary Figures

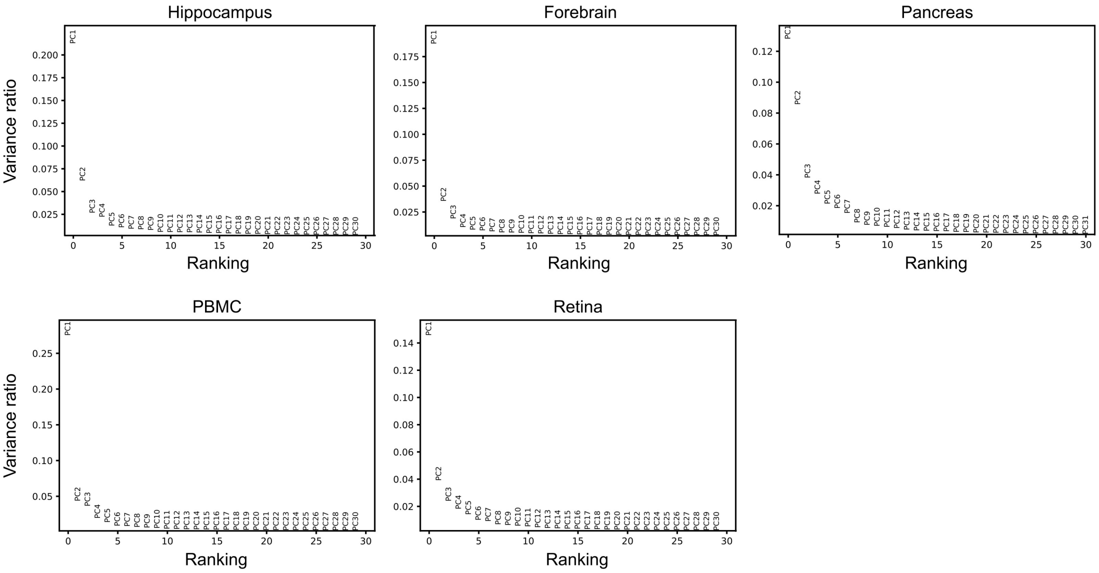

**Supplementary Figure 1: Low-rank structure of latent time.** PCA variance ratio of gene-cell specific latent time as inferred by the EM model.

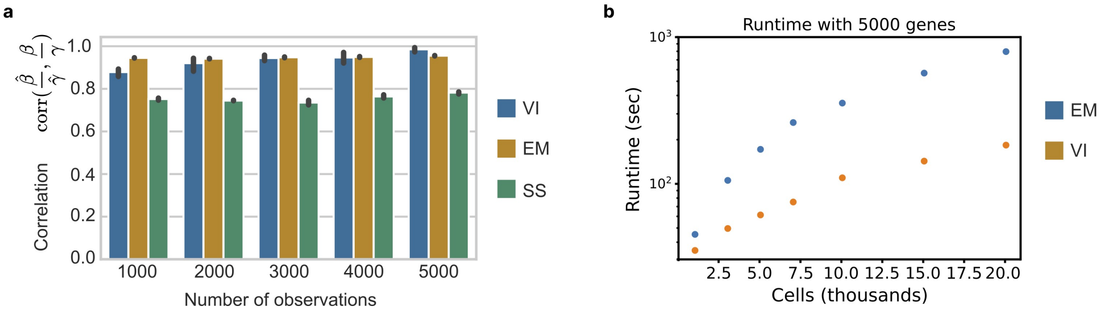

**Supplementary Figure 2: Benchmarking veloVI a.** Correlation between the estimated ratio of splicing and degradation rates, and ground truth on simulated data using veloVI (VI, blue), the *EM model* (EM, orange), steady-state model (SS, green). For each number of observations, 10 datasets were generated. **b.** Runtime comparison between veloVI (VI, orange) and the *EM model* (EM, blue). The *EM model* was run on a Intel(R) Core(TM) i9-10900K CPU @ 3.70GHz with 8 cores, veloVI on a Nvidia RTX3090 GPU.

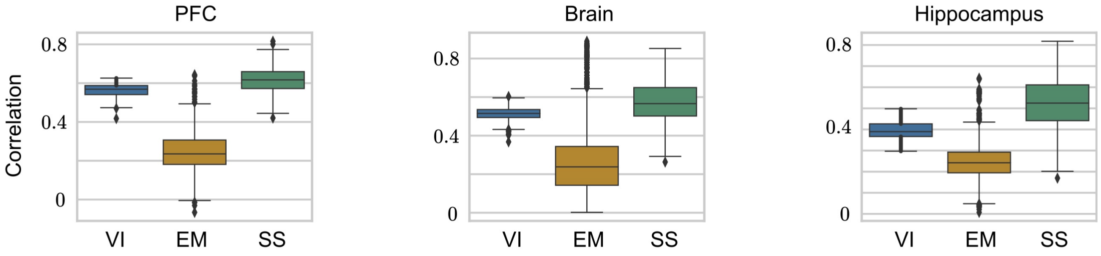

**Supplementary Figure 3: Preprocessing stability of inference methods.** Correlation of velocity estimates for datasets of prefrontal cortex (PFC) (left), 21-22 months old mouse brains (middle), and hippocampus (right). Unspliced and spliced counts are quantified with different algorithms [7, 8, 9, 10, 11, 12]. Velocities are estimated by veloVI (VI, blue), the EM model (EM, orange), and the steady-state model (SS, green).

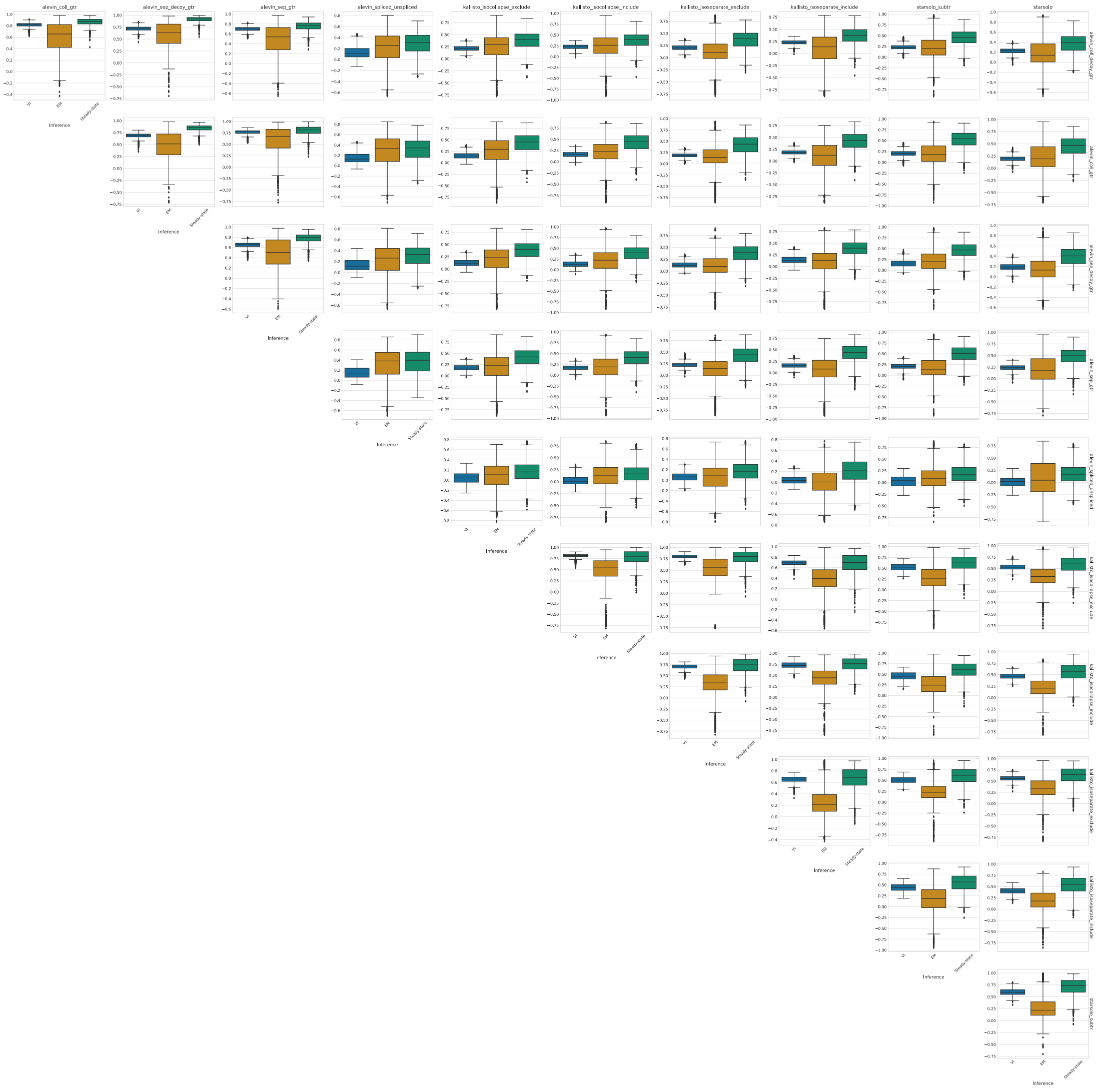

**Supplementary Figure 4: Preprocessing robustness for the dentate gyrus dataset.** Pair-wise correlations of velocity between pre-processing protocols. Velocities are inferred using veloVI, the EM model, or the steady-state model. For each pair of quantification algorithms and inference method, the correlation between the two inferred velocities for a cell are correlated.

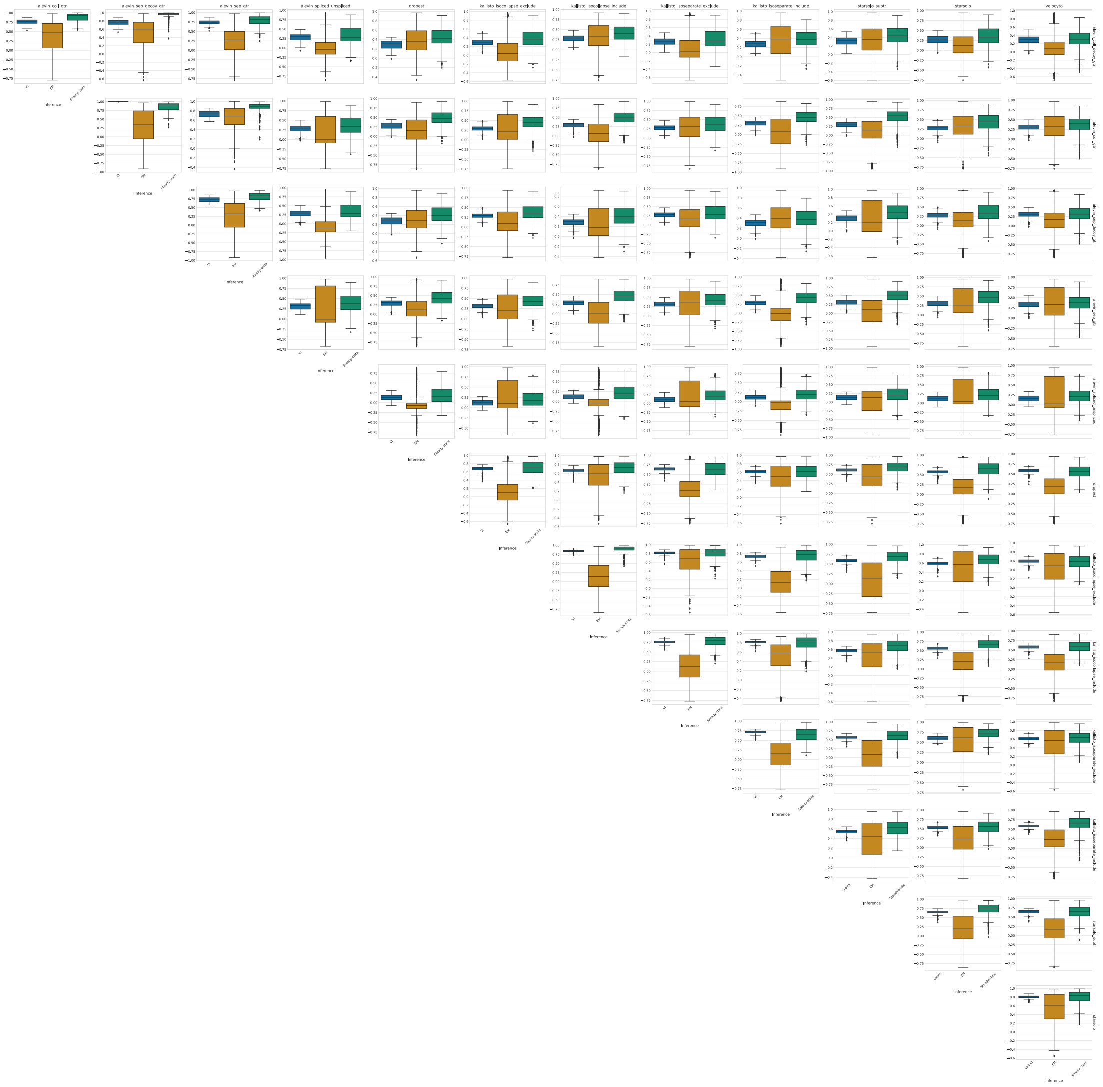

**Supplementary Figure 5: Preprocessing robustness for the old brain dataset.** Pair-wise correlations of velocity between pre-processing protocols. Velocities are inferred using veloVI, the EM model, or the steady-state model. For each pair of quantification algorithms and inference method, the correlation between the two inferred velocities for a cell are correlated.

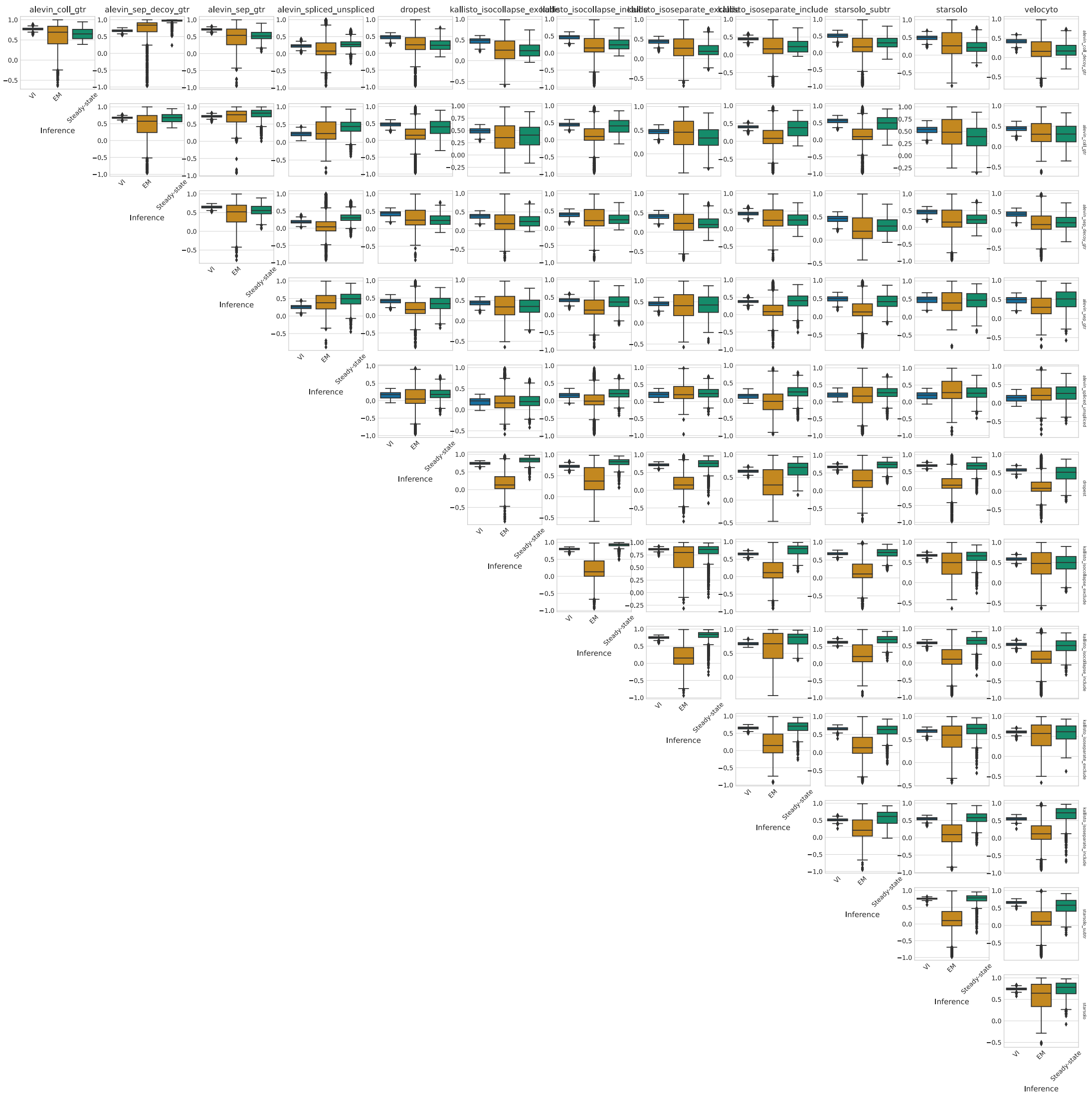

**Supplementary Figure 6: Preprocessing robustness for the pancreas dataset.** Pair-wise correlations of velocity between pre-processing protocols. Velocities are inferred using veloVI, the EM model, or the steady-state model. For each pair of quantification algorithms and inference method, the correlation between the two inferred velocities for a cell are correlated.

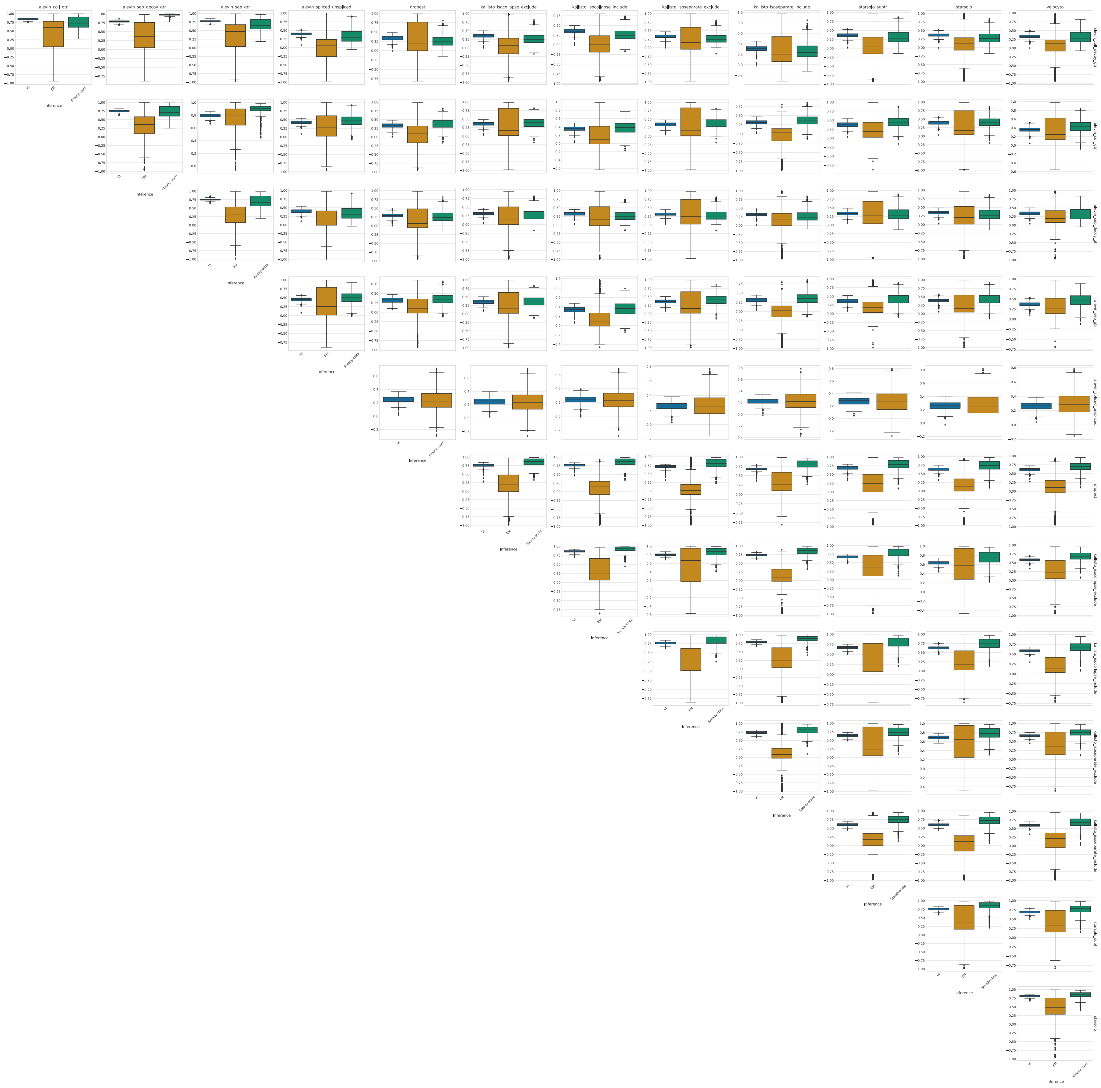

**Supplementary Figure 7: Preprocessing robustness for the PFC dataset.** Pair-wise correlations of velocity between pre-processing protocols. Velocities are inferred using veloVI, the EM model, or the steady-state model. For each pair of quantification algorithms and each inference method, the correlation between the two inferred velocities for a cell are correlated.

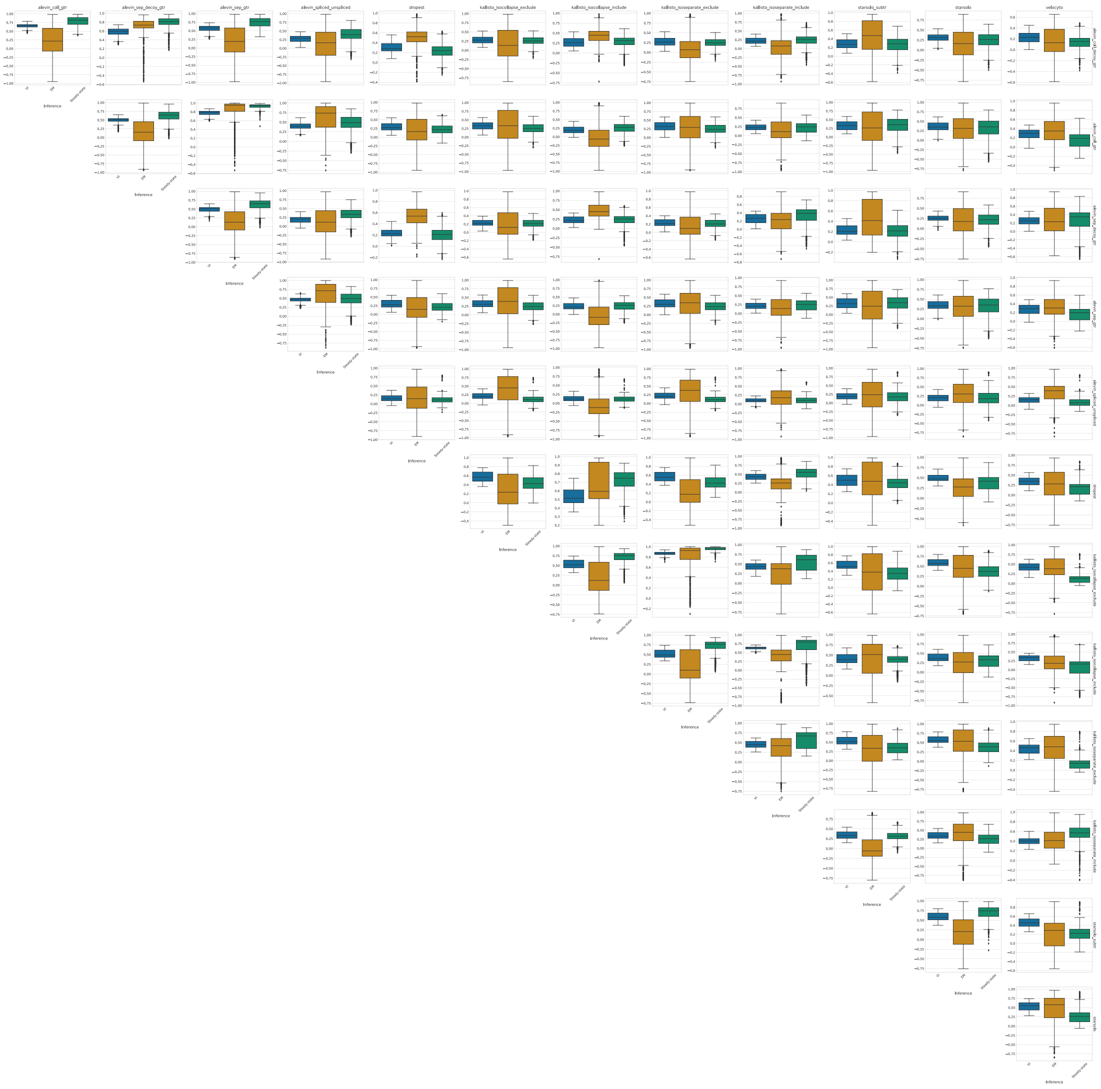

**Supplementary Figure 8: Preprocessing robustness for the spermatogenesis dataset.** Pair-wise correlations of velocity between pre-processing protocols. Velocities are inferred using veloVI, the EM model, or the steady-state model. For each pair of quantification algorithms and inference method, the correlation between the two inferred velocities for a cell are correlated.

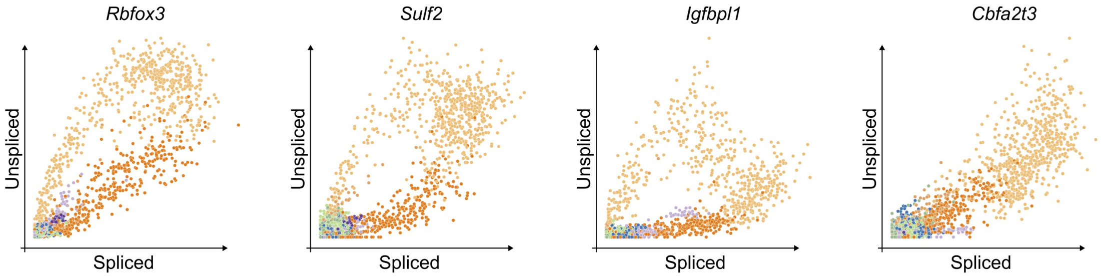

**Supplementary Figure 9: Phase portraits in pancreas endocrinogenesis.** Phase portraits of *Rbfox3*, *Sulf2*, *Igfbpl1*, and *Cbfa2t3*. Each cell is colored by its cell type. The color code is according to Figure 1 and Figure 2.

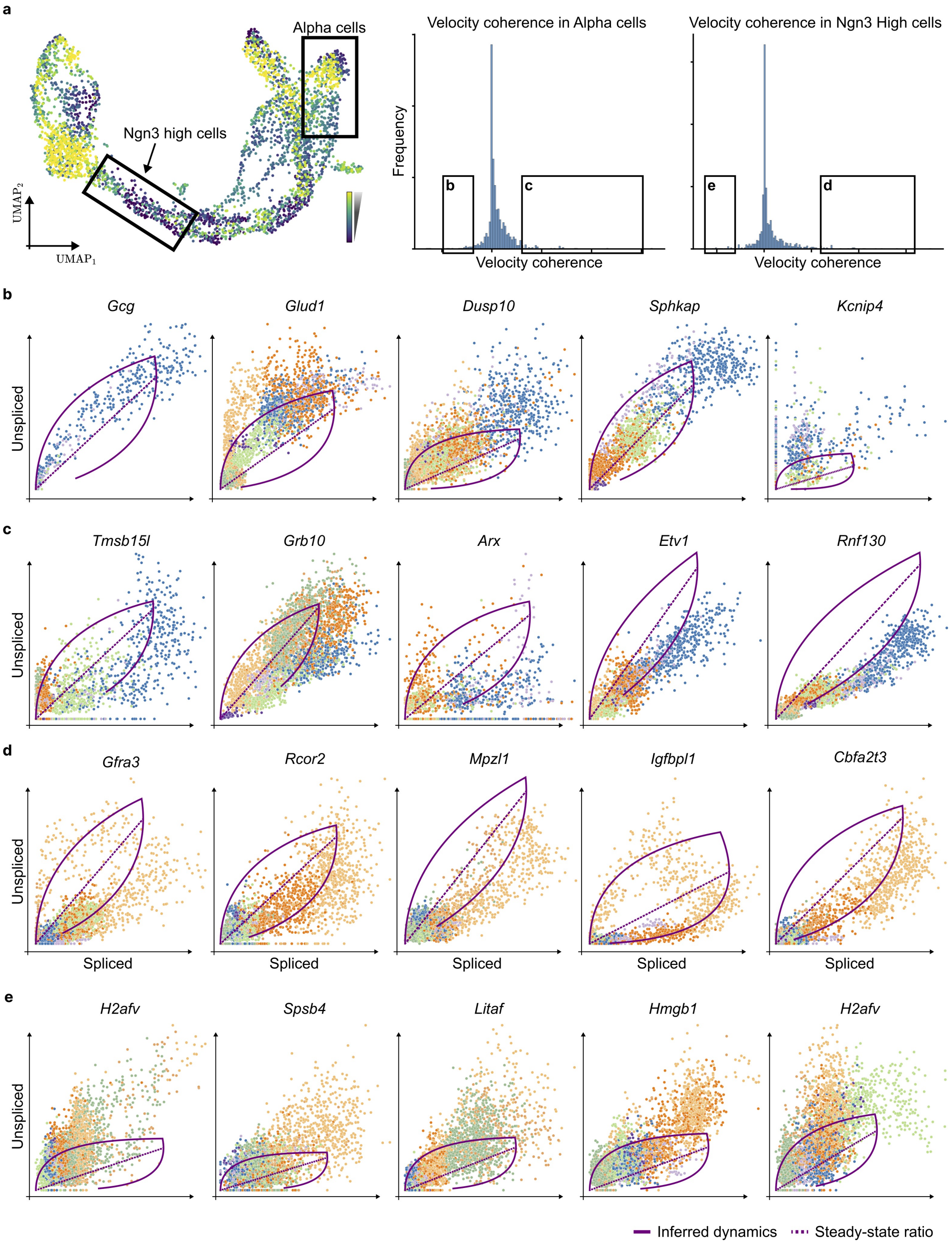

**Supplementary Figure 10: Gene analysis based on extrinsic uncertainty. a.** UMAP embedding of the Pancreas dataset colored by extrinsic uncertainty (left); The velocity coherence score across all genes for *Alpha* and Ngn3-high cells (right). **b, c.** Genes with the lowest/highest velocity coherence in *Alpha* cells, respectively. **c, d.** Genes with the lowest/highest velocity coherence in *Ngn3-high* cells, respectively.

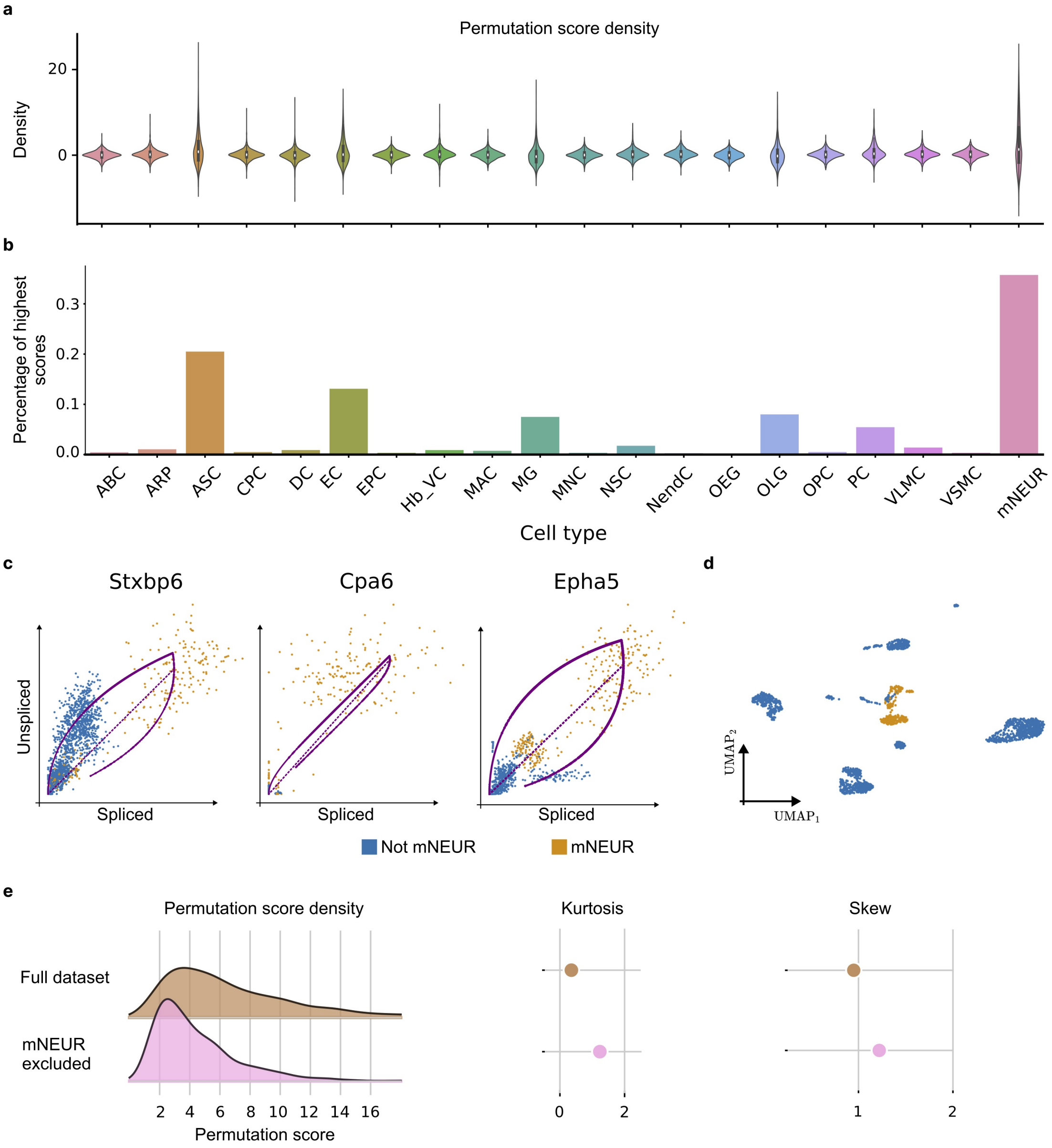

**Supplementary Figure 11: Permutation score analysis of old mouse brain. a.** Density of permutation score per cell type: arachnoid barrier cells (ABC), astrocyte-restricted precursors (ARP), astrocytes (ASC), choroid plexus epithelial cells (CPC), dendritic cells (DC), endothelial cells (EC), ependymocytes (EPC), hemoglobin-expressing vascular cells (Hb-VC), macrophages (MAC), microglia (MG), monocytes (MNC), neural stem cells (NSC), neuroendocrine cells (NendC), olfactory ensheathing glia (OEG), oligodendrocytes (OLG), oligodendrocyte precursor cells (OPC), pericytes (PC), vascular and leptomeningeal cells (VLMC), vascular smooth muscle cells (VSMC), mature neurons (mNEUR). **b.** Percentage of cell types scoring assigned the highest permutation socre for a given gene. **c.** Genes assigned the highest permutation score. **d.** Permutation score densities (left), and their kurtosis and skew when using the full dataset (brown) compared to excluding mature neurons (mNEUR).

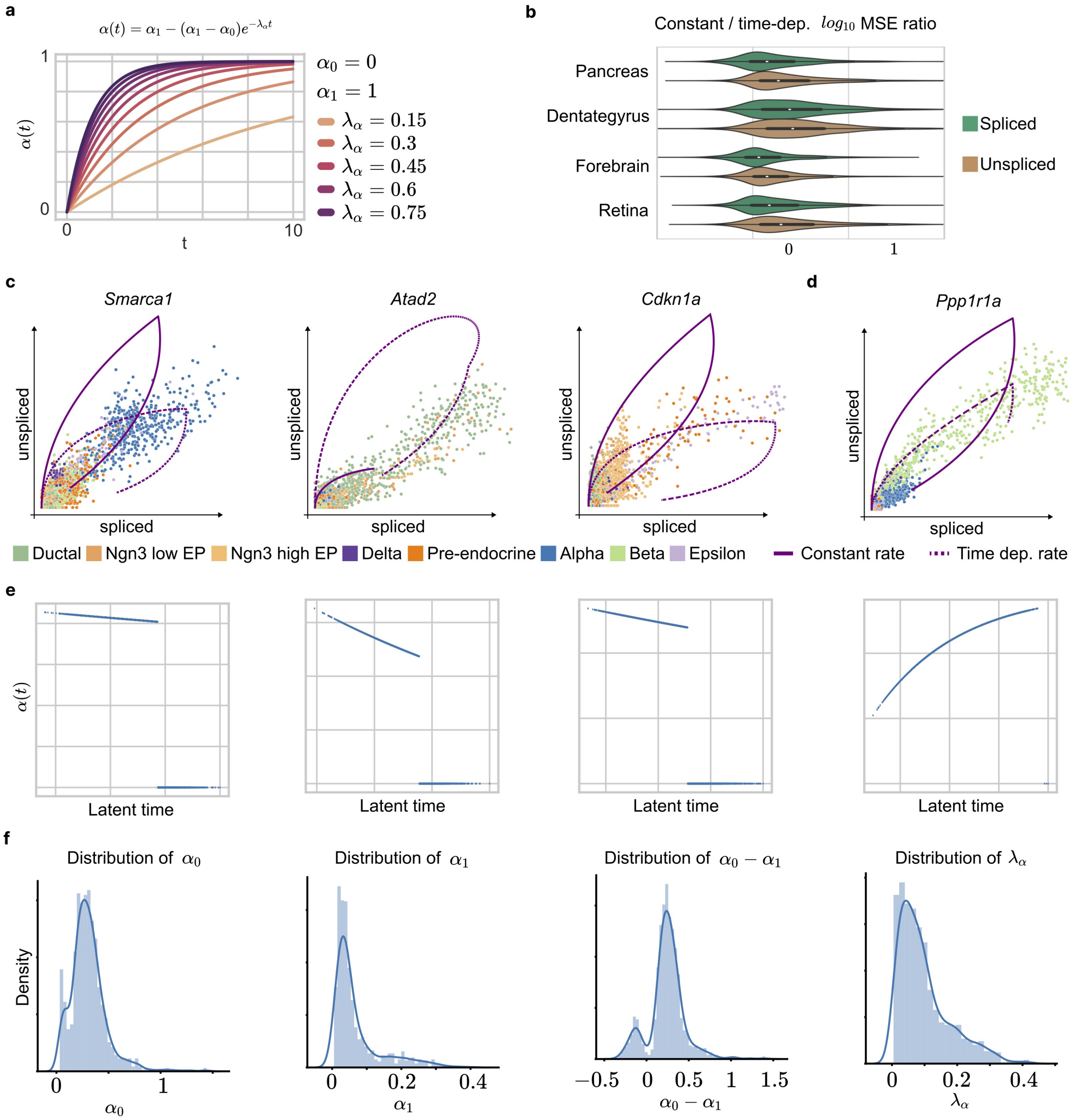

**Supplementary Figure 12: Time-dependent transcription rate. a.** Time-dependent transcription rate for different sets of parameter values. **b.** Log10 MSE ratio of models with constant and time-dependent transcription rate in the case of the pancreas, dentate gyrus, forebrain, and retina. **c.** Gene phase portrait with inferred dynamics with a constant (solid line) or time-dependent transcription rate (dashed line). Corresponding time-dependent transcription rates are shown in terms of the inferred latent time. **d.** Distribution of inferred parameters of time-dependent transcription rate.

### Supplementary Note 1

#### PBMC case study

Here, we focus on applying veloVI to a dataset of peripheral blood mononuclear cells [13, 14] (Supplementary Figure 13a). This public dataset from 10x Genomics was processed with Kallisto Bustools [9] and automatically annotated via totalVI [15] using the Seurat v3 CITE-seq PBMC dataset [13, 14] as a reference. As the dataset contains fully mature cell types, we expect the cells to be in steady-state with respect to RNA metabolism. Hence, we posit that RNA velocity cannot be used to gain insight into cell type transitions in this dataset.

After estimating velocities with veloVI and visualizing using popular techniques (Supplementary Figure 13a), we quantified the corresponding extrinsic uncertainties. The overall increased extrinsic uncertainty in the cluster labelled as “other”, as well as the substantial distribution mass of every other cell type away from the origin suggests that an analysis based RNA velocity is not suitable (Supplementary Figure 13b, c).

To assess the coherence of velocity estimates, we can study our proposed velocity coherence score. The velocity coherence in monocytes, B, and natural killer cells is close to symmetric around zero (Supplementary Figure 13d). Consequently, the inferred models include (approximately) equal number of cases in which the inferred velocity and empirical displacement of a cell agree and disagree. This result also undermines the use of RNA velocity on this dataset as there is no clear set of genes that agrees with the consensus directionality.

Finally, we calculated the permutation score to identify genes sensitive to changes in the abundance of unspliced and spliced mRNA. In the case of genes such as *TNFIAP6* and *CTSL*, the cell-type-specific permutation score is low (Supplementary Figure 13e). Consequently, these genes likely add noise to the directionality as they do not provide a signal displaying transient dynamics. This metric-based decision is confirmed by the corresponding phase portrait themselves as they do not exhibit the required (partial) almond shape translating, under the given model assumptions, to induction and repression states (Supplementary Figure 13e).

Conversely to genes ill-suited for RNA velocity analysis, we can focus on candidate genes scoring a high permutation score. The genes assigned a high permutation score across cell types included *EPHB1* and *CLEC4C*. However, the high permutation score is solely observed in dendritic cells (Supplementary Figure 13e). Again, studying the corresponding phase portraits, we can conclude that the score is likely the result of dendritic cells forming an outlier cluster as the phase portraits show discontinuity.

Taking all observations and metrics into consideration, we can conclude that caution is warranted if the dataset of PBMCs were to be analyzed using RNA velocity. This conclusion aligns with the biological ground truth that these cell types are in steady-state.

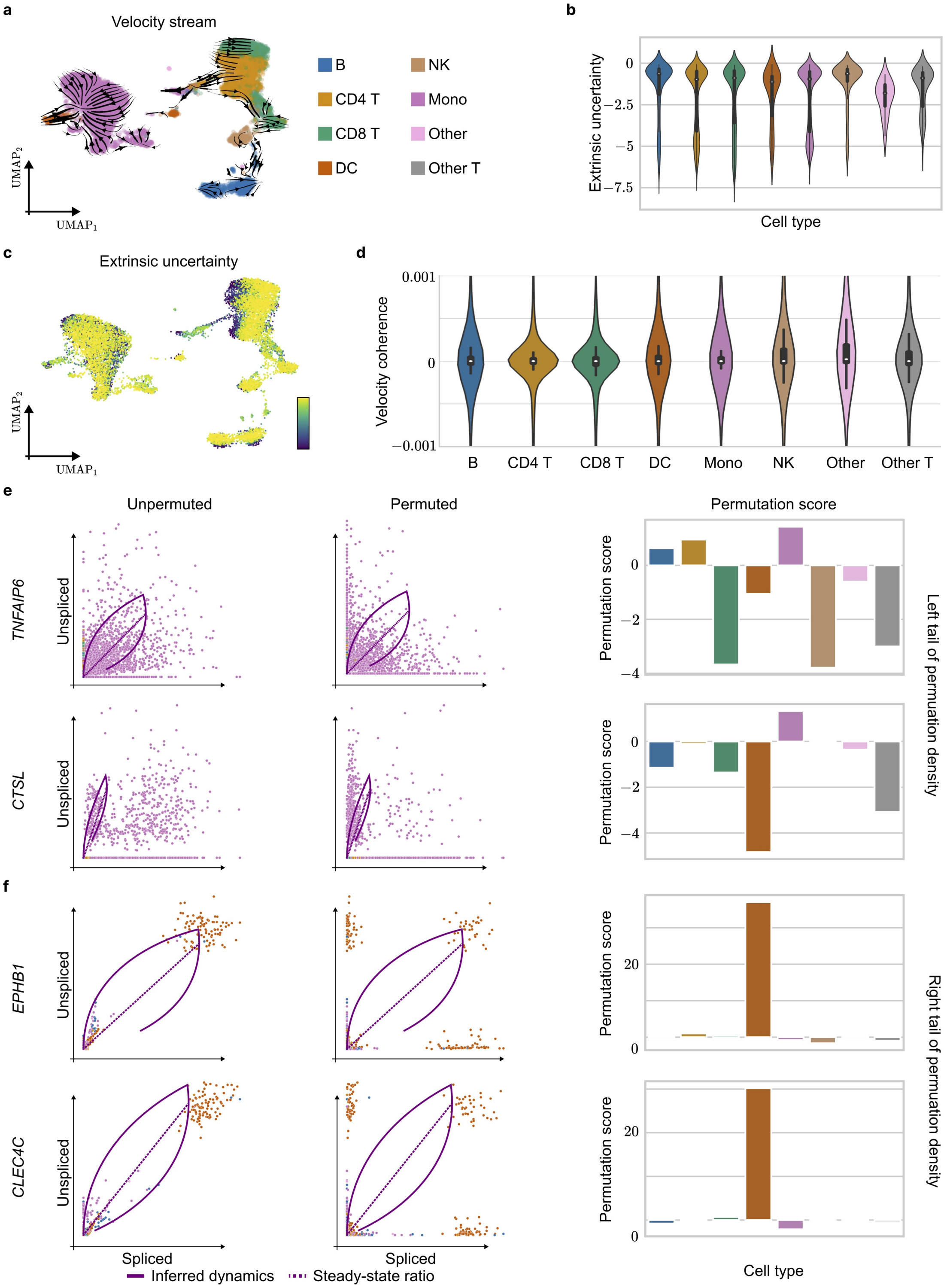

**Supplementary Figure 13: Analysis of peripheral blood mononuclear cells. a.** Two-dimension UMAP projection of PBMC dataset consisting of approximately 12,000 cells with inferred velocity stream projection. Clusters are colored by cell type (dendritic cells (DC), monocytes (Mono), natural killer (NK)). **b.** The corresponding extrinsic uncertainty resolved each cell type. Colors are the same as in panel a. **c.** The two-dimensional UMAP projection colored by the extrinsic uncertainty.**d.** The velocity coherence in each cell type. **e.** The phase portrait of the real., unpermuated data (left), permuted data (middle), and result cell-type specific permutation score. Results are shown for Genes *TNFAIP6* (top) and *CTSL* (bottom). Colors are according to cell type in panel a. **f.** Same as panel e but for *EPHB1* (top) and *CLEC4C* (bottom).

### Supplementary Note 2

#### Dentate gyrus case study

The analysis of data using RNA velocity currently usually consists in inferring velocities and projecting them onto a low-dimensional representation of the data (e.g., UMAP [17]). To highlight and showcase how our proposed veloVI method can both infer RNA velocity and aid in understanding its applicability and result, here, we conduct a case study on a dataset of dentate gyrus neurogenesis [16]. This dataset is expected to be a positive control with putative transient dynamics.

As a first step, velocities are inferred using veloVI and the corresponding stream projected onto a two-dimensional UMAP embedding of the data (Supplementary Figure 14a). Here, we observe a flow from granule mature to immature cells which is putatively incorrect according to biological ground truth (granule immature to mature cells). While the intrinsic and extrinsic uncertainties are lower for neuroblast and granule immature cells, both uncertainties are elevated in the cluster of granule mature cells (Supplementary Figure 14b, c).

In addition, the velocity coherence for a given cell type can be quantified. In case of the granule mature cells, the metric is, to a majority, positive (Supplementary Figure 14d). The positivity, thus, shows that both velocity and empirical displacement under the induced transition matrix agree. However, as the mean of the distribution is close to zero, and both the extrinsic and intrinsic uncertainties are high, velocity estimates are most likely not robustly estimated.

Similarly to the granule mature cells, we observe increased uncertainties in the clusters of endothelial, oligodendrocyte precursor (OP), myelinating oligodendrocytes (OL), and microglia cells. Additionally, the velocity coherence in these cell types is mostly zero and its distribution symmetric around the origin (Supplementary Figure 14d). Together with the fact that these cell types form distinct, disconnected clusters, we can conclude that these cell types may need to be excluded from the RNA velocity analysis. One reason to exclude these cell types is that their dynamics are distinct from the remaining dataset, or corresponding transient cell populations have not been observed. In phase portraits these cases may manifest themselves as trajectories deviating from the expected almond shape or outlier clusters.

Another tool to assess the applicability of RNA velocity to the given dataset as a whole, as well as individual genes is the permutation score. Genes scoring a large permutation score across different cell types are likely to show transient dynamics. Studying the distribution of the maximum permutation score over cell types shows that granule immature and neuroblast cells are most sensitive to permutation (Supplementary Figure 14e). This result suggests that the two populations are transient which aligns with the underlying known dynamics in dentate gyrus neurogenesis.

At the level of a single gene, the permutation score reveals that in *Tmsb10*, for example, four cell types (neuroblast, granule immature, endothelial, and GABA) are sensitive to permutation (Supplementary Figure 14e). Although the fit is confounded by the endothelial cluster, it is correctly inferred for the neuroblast to granule immature lineage. Similarly, as endothelial cells are not contained in the neuroblast to granule immature lineage, this result also shows that *Tmsb10* contains multiple kinetics. Consequently, these observations show that a single gene-specific model is inappropriate, or that only granule lineage cell types should be considered for this gene.

Genes inappropriate for RNA velocity analysis can be identified similarly based on the permutation score. The low permutation scores for each cell type in *Eed*, for example, show that it should not be considered RNA velocity analysis in the first place (Supplementary Figure 14f). The low cell-type-specific scores stem from the non-transient and noisy nature of the unspliced and spliced abundance. Consequently, they do not reflect the expected almond shape given the model assumptions, and yield the parameter inference non-robust.

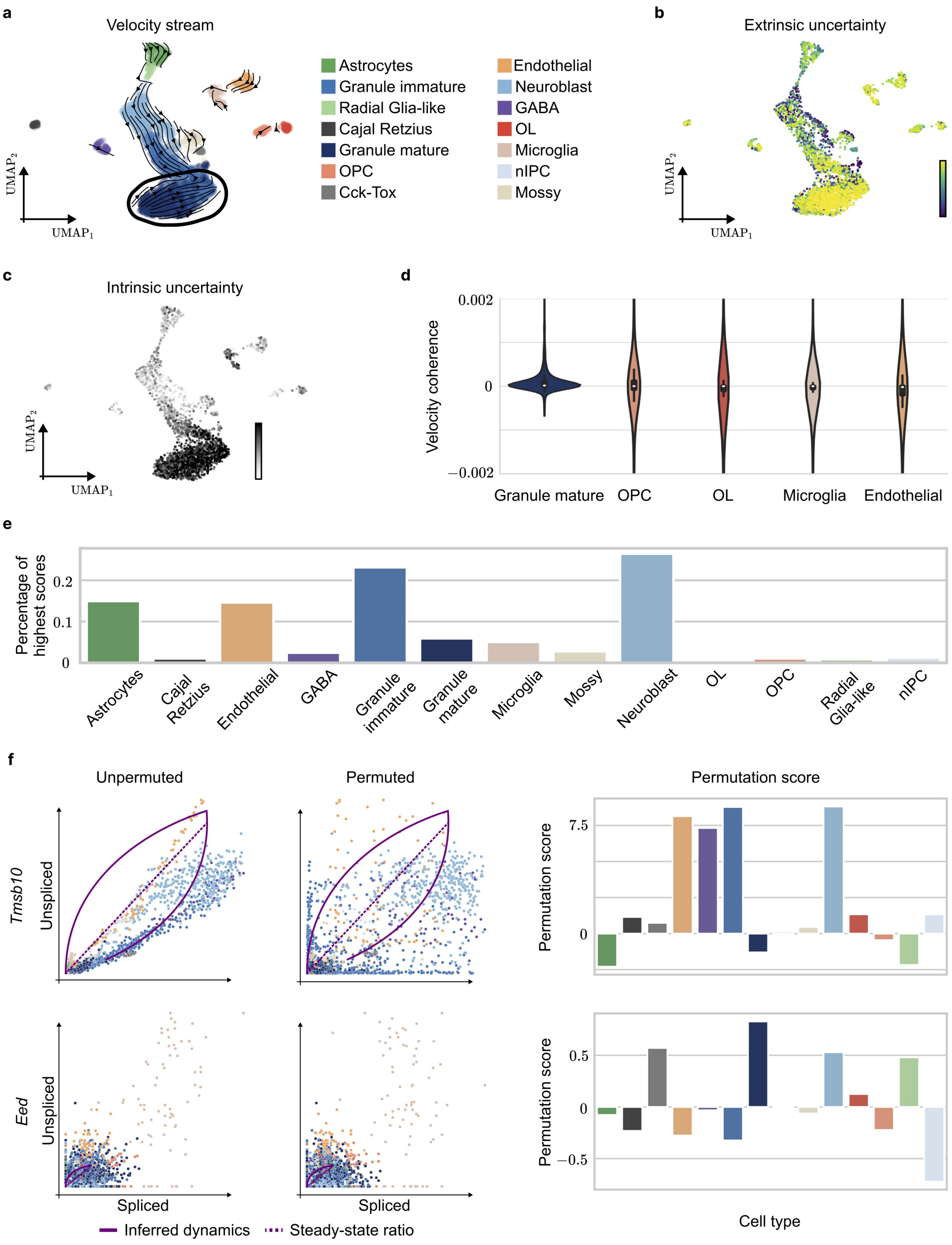

**Supplementary Figure 14: Analysis of dentate gyrus. a.** Two-dimensional UMAP representation of a dentate gyrus dataset containing 2930 cells. The corresponding velocity stream is projected onto the embedding and represented by black arrows. Each cell is colored by their cell type (oligodendrocyte progenitor cells (OPC), oligodendrocyte (OL)) according to the original work [16]. **b.** UMAP emebdding colored by extrinsic, **c.** and intrinsic uncertainty. **d.** The density of the corresponding velocity coherence is shown for granule mature cells, OPC, OL, microglia, and endothelial cells. **e.** The percentage of each cell type scoring the highest permutation score. **f.** The phase portraits of the unpermuted (left), permuted (middle) and resulting permutation score per cell type are given for *Tmsb10* and *Eed*. Coloring corresponds to cell type as defined in panel a.

### Supplementary Note 3

#### Related work

New approaches have been developed to blend RNA velocity with deep-learning-based representation learning. Here we briefly describe each approach and then relate them to the capabilities of veloVI.

**VeloAE** VeloAE [18] leverages an autoencoder framework to learn a representation of both spliced and unspliced data that can be used to estimate transcriptional dynamics. This autoencoder makes use of a graph convolutional network (GCN) module to smooth cell representations over a graph induced by standard scRNA-seq methods (e.g., principcal components analysis followed by approximate nearest neighbors) and an attention mechnaism for its decoder. RNA velocity is defined in the cell representation space of the model and and as a result has no mechanistic interpretation nor a directly interpretable link to genes.

**DeepVelo (GCN-based)** DeepVelo [19] seeks to generalize RNA velocity to multi-lineage systems with cell-specific kinetics. To achieve this, DeepVelo leverages GCNs to encode spliced and unspliced abundance for a cell while aggregating over spliced-abundance-induced neighbourhood graph. In the decoding phase, DeepVelo predicts cell-specific $\alpha$, $\beta$, and $\gamma$ parameters and uses the mechanistic definition of velocity to compute a cell-specific velocity. This velocity is trained to be predictive of future cell states under a first-order approximation and future cell states are restricted to nearest neighbors. This assumption is referred to as the "continuity assumption". As cell neighborhoods will contain future and past cell states under this assumption, DeepVelo uses adds a loss term that enforces velocity to be correlated (resp., anticorrelated) with unspliced (resp., spliced) abundance. While DeepVelo leverages ideas from RNA velocity, it does not explicitly model time and thus makes use of standard pseudotime procedures based on its outputs. A similar mechanism of using cells from the training data as a proxy to infer future states of a cell has also been proposed in parallel by Marot-Lassauzaie et al.[20].

**DeepVelo (Neural-ODE-based)** DeepVelo [21] combines representation learning with neural-network-based ordinary differential equations to learn velocity fields. The model leverages denoising variational autoencoders (VAE) to learn a mapping between the spliced abundance and the velocity estimated with scVelo. They further leverage the VAE within external black-box ordinary differential equation solvers to simulate past or future states of each cell.

**DeepCycle** In contrast to previous approaches, DeepCycle [22] does not estimate RNA velocity parameters. The method exploits the concept of RNA velocity and its connection to cell-cycle to infer a 1-dimensional latent variable representing the cell-cycle phase using an autoencoder framework.

**VeloVAE** Upon finalizing this manuscript, we came across two independent manuscripts describing VeloVAE [23, 24]. VeloVAE uses a variational autoencoder framework to estimate kinetic parameters and learns a posterior distribution over a cell-specific latent time. The model likelihood makes use of the solution of ordinary differential equations describing RNA metabolism. VeloVAE posits a cell-gene specific transcriptional rate, which is the function of a low-dimensional latent variable (capturing cell state). Uncertainty in the VeloVAE model is quantified by a coefficient of variation on the low-dimensional latent variable.

**Relation to veloVI** Our approach veloVI exploits the variational inference (VI) [3] framework to infer each cell's latent time and transcriptional latent state via a local low-dimensional latent variable. Compared to the existing methods, veloVI is the only method that directly estimates RNA velocity at the level of a cell and gene via a biophysical model of transcriptional dynamics *and* leverages uncertainty in its estimate of velocity in downstream applications. The closest approaches directly estimating transcriptional parameters $\alpha$, $\beta$, and $\gamma$ are DeepVelo (GCN-based) [19] and VeloVAE [23, 24]. Yet, DeepVelo (GCN-based) learns these parameters such that they produce velocities that conform to the model's "continuity" assumption, as well as the notion that velocity should be correlated with unspliced abundance while anticorrelated with spliced abundance; therefore, it only retains a loose connection to transcriptional dynamics.

At a high level, VeloVAE is conceptually similar to veloVI – both approaches make use of the solved differential equations in the model likelihood and manifest as variational autoencoders. For both models, velocity is computed as a statistical functional of the variational posterior. However, there are key differences.

Compared to VeloVAE, veloVI offers unique features to aid in RNA velocity analysis: Intrinsic and extrinsic uncertainty of the estimated velocity, velocity coherence, and the permutation score. Quantifying uncertainty at the level of RNA velocity (cell-gene-specific, and then aggregated) allows assessing regions of the transcriptomic manifold where RNA velocity is either well supported (low uncertainty), or further investigation is needed (high uncertainty). This information is complemented by the permutation score that helps identify viable genes and transient cell types. Consequently, the parameter inference can be quantified beyond a visual, low-dimensional representation. The permutation score based analysis, allows, for example, to correctly identify a dataset of peripheral blood mononuclear cells as inappropriate for RNA velocity analysis (Supplementary Note 1).

veloVI considers uncertainty over transcription states (e.g., induction and repression) for parameter inference. Contrastingly, VeloVAE uses a cell-gene specific transcription rate, as well as a cell-specific latent time. While VeloVAE preprint [23] includes model uncertainty, the uncertainty is at the level of cells via the cell-specific latent representation (analog to $z_n$ in veloVI). This cell-specific representation has a complex non-linear relationship to the estimated velocity (via a neural network).

While veloVI offers tools to assess if a given dataset is suitable for RNA velocity analysis, VeloVAE does not. Indeed, false positives persist even with VeloVAE's improvements. For example in Figure 4 of ref. [23], spurious transitions appear on the velocity stream plot in a dataset of human bone marrow mononuclear cells going from Memory B to Naive B cell types, as well as going from CD8 memory to NK cell types (both with low VeloVAE cell state uncertainty; Supplementary Figure 5e of ref. [23]).

### Supplementary Note 4

#### Splicing kinetics with time-dependent rates

To highlight veloVI's extensibility w.r.t. model choice, we consider the time dependent transcription rate

$$\alpha^{(k)}(t) = \begin{cases} \alpha_1 - (\alpha_1 - \alpha_0)e^{-\lambda_\alpha t}, & k \in \{1, 2\}, \\ 0, & k \in \{3, 4\}, \end{cases}$$

with parameters $\alpha_0, \alpha_1, \lambda_\alpha \in \mathbb{R}^+$, and $k$ indicating the transcriptional state ($k = 1$ induction, $k = 2$ induction steady-state, $k = 3$ repression, $k = 4$ repression steady-state). The system of differential equations describing the process of splicing stays otherwise unchanged and is, thus, given by

$$\begin{aligned} \dot{u} &= \alpha^{(k)}(t) - \beta u \\ \dot{s} &= \beta u - \gamma s. \end{aligned} \tag{33}$$

Consequently, it is of the general form

$$\dot{x} = Ax + g(t), \tag{34}$$

with dependent variable $x$, system matrix $A$, inhomogeneity $g(t)$, and solution

$$x(t) = x_0 e^{A(t-t_0)} + e^{At} \int_{t_0}^{t} e^{-As} g(s) ds. \tag{35}$$

As the abundance of unspliced mRNA is modelled independently of its spliced counterpart, its solution of (33) can be found directly. Comparing (33) with (34) and (35), we find $x = u$, $A = -\beta$, $g(t) = \alpha^{(k)}(t)$. Consequently, the abundance of unspliced mRNA at time $t$ is given by

$$\begin{aligned} u(t) = \quad & u_0^{(k)} e^{-\beta\tau^{(k)}} + \alpha_1^{(k)} e^{-\beta t} \int_{t_0^{(k)}}^{t} e^{\beta s} ds - (\alpha_1^{(k)} - \alpha_0^{(k)}) e^{-\beta t} \int_{t_0^{(k)}}^{t} e^{\beta s} e^{-\lambda_\alpha^{(k)} s} ds \\ = \quad & u_0^{(k)} e^{-\beta\tau^{(k)}} + \frac{\alpha_1^{(k)}}{\beta} \left(1 - e^{-\beta\tau^{(k)}}\right) \\ & - \frac{\alpha_1^{(k)} - \alpha_0^{(k)}}{\beta - \lambda_\alpha^{(k)}} e^{-\lambda_\alpha^{(k)} t_0^{(k)}} \left(e^{-\lambda_\alpha^{(k)} \tau^{(k)}} - e^{-\beta\tau^{(k)}}\right), \end{aligned} \tag{36}$$

with state-dependent initial time $t_0^{(k)}$, $\tau^{(k)} = t - t_0^{(k)}$, and $u_0^{(k)} = u(t_0^{(k)})$.

Similarly, this allows solving for $s(t)$, with $x = s$, $A = -\gamma$, $g(t) = \beta u(t)$. Applying solution formula (35), the abundance of spliced mRNA at time $t$ is given by

$$\begin{aligned} s(t) = \quad & s_0^{(k)} e^{-\gamma\tau^{(k)}} + e^{-\gamma t} \int_{t_0^{(k)}}^{t} e^{\gamma t'} \beta u(t') dt' \\ = \quad & s_0^{(k)} e^{-\gamma\tau^{(k)}} + \frac{\alpha_1^{(k)}}{\gamma} \left(1 - e^{-\gamma\tau^{(k)}}\right) + \frac{\alpha_1^{(k)} - \beta u_0^{(k)}}{\gamma - \beta} \left(e^{-\gamma\tau^{(k)}} - e^{-\beta\tau^{(k)}}\right) \\ & - \frac{\beta(\alpha_1^{(k)} - \alpha_0^{(k)})}{(\beta - \lambda_\alpha^{(k)})(\gamma - \lambda_\alpha^{(k)})} e^{-\lambda_\alpha^{(k)} t_0^{(k)}} \left(e^{-\lambda_\alpha^{(k)} \tau^{(k)}} - e^{-\gamma\tau^{(k)}}\right) \\ & + \frac{\beta(\alpha_1^{(k)} - \alpha_0^{(k)})}{(\beta - \lambda_\alpha^{(k)})(\gamma - \beta)} e^{-\lambda_\alpha^{(k)} t_0^{(k)}} \left(e^{-\beta\tau^{(k)}} - e^{-\gamma\tau^{(k)}}\right), \end{aligned} \tag{37}$$

### Supplementary Tables

| Dataset | Reference | Organism | Number of observations |
|---|---|---|---|
| Pancreatic endocrinogenesis | [25] | Mouse | 3,696 |
| Spermatogenesis | [26, 27] | Mouse | 1,829 |
| Hippocampus | [7] | Mouse | 18,213 |
| Forebrain | [7] | Human | 1,720 |
| Retina | [28, 29] | Mouse | 2,726 |
| Brain | [30, 27] | Mouse | 1,823 |
| Prefrontal cortex | [31, 27] | Mouse | 1,267 |
| PBMC | [13] | Human | 11,950 |
| Dentate gyrus neurogenesis | [16] | Mouse | 2,930 |
| Retina (Runtime analysis only) | [32] | Mouse | 113,909 |

**Supplementary Table 1: Overview of datasets used in this manuscript.**
